# Stressor identity shapes plasma proteomic and metabolic responses in humans

**DOI:** 10.64898/2026.07.17.739102

**Authors:** T Ebert, T Bajaj, J Stepan, B Herhaus, J Eberz, W Bi, S Kallabis, E Junglas, M Lennarz, DE Heinz, A Brahmer, MC Sokn, S Frei, S Bulut, D Stanivuk, J Strupp, H Hohenthanner, P Bajaj, M Bergmann, NC Braun, T Fischer, E Halbe, M Huppertz, B Jurek, H Jaksch-Bogensperger, S Kaspar-Schrödl, V Kempf, M Kölle, E Kratz, A Kudaliyanage, M Laakmann, K Lammertz, C Ledschbor-Frahnert, B Leeners, A Lindner, S Lux, S Mackert, S Maier, B Mayr, Y Mecdad, G Mohsen, C Niemeyer, K Prominski, A Rabaszowski, H Rohner, M Saleh, M Sareban, MV Schmidt, J Tumusiime, E Tuncöz, A Zimmermann, H Schneider, A Philipsen, MB Müller, B Steigenberger, J Cox, K Gapp, K Petrowski, EM Krämer-Albers, N Paczia, F Meissner, NC Gassen

## Abstract

Acute stress triggers rapid physiological adaptations, yet how distinct stress modalities shape the human circulating molecular landscape remains unclear. We performed longitudinal deep plasma proteomics and targeted metabolomics across three paradigms: psychological stress, controlled physical stress, and combined psychological–physical stress induced by bungee jumping. Despite robust hypothalamic–pituitary–adrenal axis activation, psychological stress induced minimal plasma proteomic and metabolic changes, whereas physical stress triggered rapid, coordinated proteome remodeling accompanied by stressor-specific metabolic adaptations. Combined stress elicited the strongest and most persistent proteomic response, defining a graded molecular signature across stress modalities. Cross-paradigm integration identified a stress-associated protein core enriched for immune granule effectors, RNA-binding proteins, and vesicle-linked signatures, suggesting regulated extracellular release. Consistently, extracellular vesicle profiling in the mixed-stress paradigm supported vesicle-associated secretion, while functional experiments indicated that glucocorticoid signaling alone is insufficient to trigger release. Together, these findings reveal stressor identity as a key determinant of the human circulating molecular response.

## Introduction

The acute stress response is a fundamental physiological adaptive process that enables organisms to cope with environmental challenges. Triggered by perceived threats ranging from psychological and social stressors to intense physical demands, acute stress rapidly mobilizes neuroendocrine, metabolic, and immune systems to support adaptive behavioral and physiological responses (Chrousos & Gold, 1992). Under healthy conditions, these responses are tightly regulated and resolved once the threat subsides, allowing restoration of homeostasis. In contrast, prolonged or dysregulated stress responses contribute to the development of numerous pathological conditions, including cardiovascular disease, metabolic dysfunction, and psychiatric disorders (de Kloet & Joëls, 2023; McEwen et al., 1998).

Historically, stress research has focused on central nervous system circuits and activation of the hypothalamic–pituitary–adrenal (HPA) axis. However, growing evidence indicates that acute stress rapidly reshapes the circulating molecular environment, reflecting coordinated responses across multiple organs and physiological systems (Rohleder et al., 2019; Galluzzi et al., 2018). Plasma serves as a central integrative compartment for systemic signaling, containing endocrine mediators, immune-derived proteins, metabolic intermediates, and cell-derived vesicles that together mediate the inter-organ communication. Advances in high-resolution omics technologies have begun to reveal the complexity of these circulating responses. Plasma proteomics and metabolomics studies have identified stress-associated changes in inflammatory mediators, metabolic pathways, and acute-phase proteins in humans exposed to psychological or physiological challenges (Marsland et al., 2017; Steptoe et al., 2007; Contrepois et al., 2020). Despite these advances, a central question remains unresolved: do different forms of acute stress (e.g. physical versus social-emotional stress) engage shared systemic molecular programs, or do they elicit fundamentally distinct circulating response patterns?

This question has important implications for advancing our understanding of stress-related pathophysiology. Psychological and physical stressors differ fundamentally in their cognitive appraisal, metabolic demands, mechanical load and in their valence: whereas exercise represents a predominantly positive, adaptive stressor (eustress), psychological stress is frequently experienced as negative and uncontrollable (distress), and is strongly linked to adverse health outcomes when chronic or dysregulated (Heidt et al., 2014; Knezevic et al., 2023; Herhaus et al., 2024; Gaab et al., 2005). Despite this fundamental difference in valence, both modalities reliably activate canonical neuroendocrine pathways, particularly the HPA axis. Determining whether these stress modalities converge on common systemic pathways or instead trigger modality-specific circulating responses is essential for understanding how stress signals propagate throughout the organism. Beyond identifying stress-responsive molecules, an additional open question concerns the mechanisms by which intracellular stress signals are transmitted into the circulation. Increasing evidence suggests that non-classical secretion pathways, including extracellular vesicle (EV) release and unconventional protein secretion, play key roles in intercellular stress communication. EVs can transport proteins, RNA-binding factors, and damage-associated molecular patterns (DAMPs) between tissues, thereby propagating stress signals systemically (Tkach and Théry, 2016; van Niel et al., 2018). However, whether such mechanisms contribute to the circulating molecular signatures of acute stress in humans remains largely unexplored.

Here, we performed longitudinal deep plasma proteomics and targeted metabolomics across three complementary human stress paradigms: purely psychological stress induced by the Trier Social Stress Test (TSST), controlled physical stress induced by maximal exercise on a bicycle ergometer, and mixed psychological–physical stress induced by bungee jumping. These paradigms span a graded spectrum of stress exposure, differing in cognitive appraisal, physiological load, and emotional salience while reliably eliciting acute stress responses. Using nanoparticle-enabled plasma proteomics (Blume et al., 2020), together with targeted metabolomics, we quantified thousands of circulating molecules at multiple timepoints to resolve stressor-specific molecular trajectories. Across these paradigms, we identify a stress-associated protein core that emerges under conditions of intense physiological stress and becomes amplified during combined psychological–physical challenge. Integrating EV-profiling with cellular and in vivo perturbation experiments, we further implicate EV-mediated unconventional secretion pathways as a central mechanism linking intracellular stress states to systemic plasma remodeling.

## Results

### Psychological stress induces robust endocrine activation but minimal plasma proteomic changes

To define how distinct stress modalities shape systemic molecular responses in humans, we first examined changes in plasma proteins in response to purely psychological stress using plasma samples collected during the TSST in a cohort comprising 20 healthy controls and 20 individuals diagnosed with panic disorder (Table S1). The TSST is a highly standardized and widely used laboratory paradigm that robustly induces acute psychosocial stress through socially evaluative public speaking and mental arithmetic (Kirschbaum et al., 1993). Plasma samples were obtained at multiple timepoints before and after stress exposure, enabling longitudinal assessment of endocrine, proteomic, and metabolomic response (Fig. 1A, B). Serum cortisol measurements confirmed robust activation of the HPA axis across participants, with cortisol levels peaking shortly after stress induction and gradually returning toward baseline, without differences between patients with panic disorder and healthy controls (Fig. 1C).

**Figure 1.**
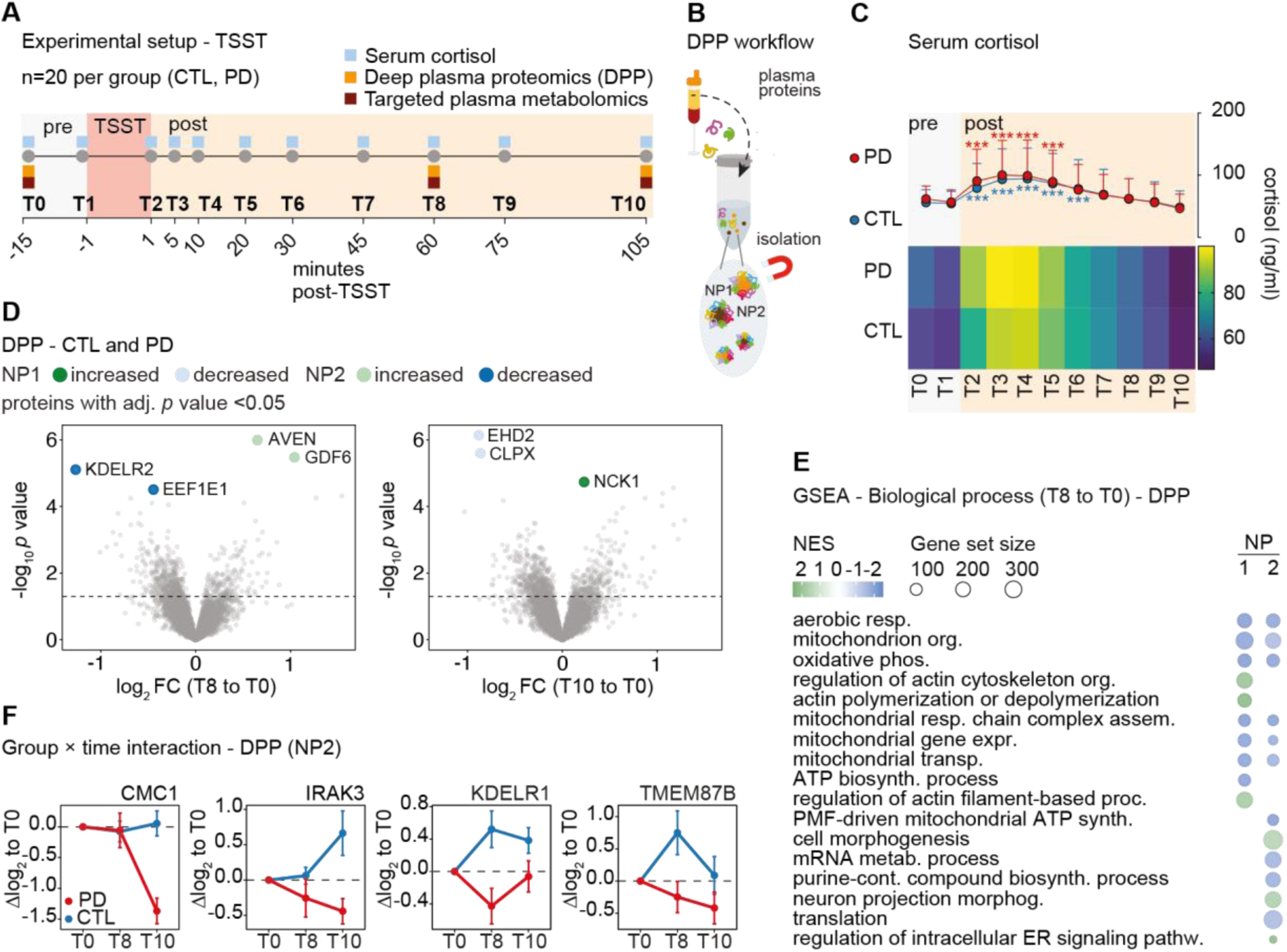
Psychological stress induces robust endocrine activation but minimal plasma proteomic remodeling. **(A)** Experimental design of the Trier Social Stress Test (TSST) cohort. Plasma samples were collected at 11 timepoints (T0–T10) spanning 15 min before the TSST procedure (T0), and up to 105 minutes post-TSST in n = 20 healthy controls (CTL) and n=20 individuals with panic disorder (PD). Serum cortisol measurements, deep plasma proteomics (DPP) and targeted plasma metabolomics were performed at indicated timepoints. **(B)** DPP workflow schematic. Plasma proteins are fractionated using nanoparticles prior to LC-MS analysis. **(C)** Serum cortisol levels (line plot, mean + SD) and heatmap of cortisol trajectories across timepoints for PD patients and CTL. Shading indicates the post-TSST period (n = 20 controls, n = 20 PD patients, Linear mixed model with post-hoc comparisons of estimated marginal means against baseline (−15 min) using Dunnett-type contrasts; Holm–Bonferroni correction for multiple comparisons. *** p < 0.001)). **(D)** Volcano plots showing differential protein abundance comparing post-TSST timepoints T8 (left) and T10 (right) to pre-stress baseline (T0) in the combined cohort (CTL and PD). (Moderated paired t-tests; Benjamini-Hochberg (BH) correction for multiple testing; adjusted p < 0.05 highlighted) **(E)** Gene Set Enrichment Analysis (GSEA) of ranked proteomic data (T8 vs. T0) for biological process gene sets, performed on the combined cohort. **(F)** Group × time interaction analysis (DPP, NP2). Line plots show log₂ fold-change relative to T0 for proteins with the lowest interaction p-values between PD and CTL groups. Data shown as mean ± SEM.

To enable sensitive detection of stress-induced changes in the circulating proteome, we implemented a novel deep plasma proteomics approach utilizing surface-functionalized superparamagnetic nanoparticles (NPs) to fractionate plasma proteins (Fig. 1B). This optimized workflow markedly enhanced our detection capability, enabling quantification of > 6000 proteins per sample after filtering across the TSST dataset (Fig. S1A-F). Each sample was fractionated with two types of nanoparticles (NP1/2, Fig. 1B) and analyzed separately via LC-MS, providing high-resolution insights into plasma protein abundance.

Despite this extensive proteomic depth, differential abundance analysis comparing post-stress time points with the immediate pre-stress baseline revealed only a few significantly regulated proteins (adjusted p < 0.05) following TSST exposure, both in the combined cohort (Fig. 1D; Table S2) and when analyzing the two groups separately (Fig. S2A-H; Table S3). Most proteins exhibited small effect sizes without evidence of coordinated or sustained regulation. Notably, several of the significantly regulated proteins were intracellular stress- and cytoskeleton-associated factors previously reported in circulating plasma proteomes, including the adaptor protein NCK1 (Fig. 1D).

Gene set enrichment analysis of ranked proteomic data revealed a selective but reproducible enrichment of mitochondrial and energy-related pathways (Fig. 1E, Fig. S3A; Table S4). In particular, terms related to aerobic respiration and oxidative phosphorylation were among the most consistently enriched gene sets following TSST exposure, suggesting distinct engagement of systemic energy metabolism.

To test whether psychological stress elicits distinct systemic molecular responses in a clinically vulnerable population, we compared plasma proteomic changes between individuals with panic disorder and healthy controls. Despite known heightened stress sensitivity in panic disorder (Abelson et al., 2007), the circulating proteomic response to stress was largely conserved across groups, with no reproducible differences detected at any post-stress timepoint (Fig. S3B-G; Table S5).

Several proteins displayed qualitatively divergent temporal trajectories between controls and patients with panic disorder (Fig. 1F; Table S6). While these differences did not reach statistical significance, the proteins with the lowest interaction p-values included regulators of innate immune regulation (IRAK3), mitochondrial function (CMC1), and ER–Golgi trafficking (KDELR1), suggesting subtle differences in cellular stress adaptation rather than exaggerated systemic plasma responses.

Together, these findings reveal a dissociation between endocrine activation and systemic plasma proteomic remodeling during acute psychological stress. Despite robust HPA axis activation, the circulating proteome remained largely stable in both healthy individuals and participants with panic disorder. Instead, the detectable signal pointed to subtle metabolic adaptation, suggesting that acute psychological stress is reflected more prominently in selective metabolic pathways rather than large-scale protein release. This relative stability of the circulating proteome during purely psychological stress provided an important reference point for interpreting responses to more intense physiological challenges examined in the subsequent stress paradigms.

### Physical stress induces coordinated plasma proteome remodeling

To define the circulating proteomic response to acute physical stress, we next profiled plasma samples from 15 recreationally active men, performing a standardized incremental cycling test to volitional exhaustion reaching their individual maximal oxygen uptake (VO_2_max, (Fig. 2A, Table S8;). Plasma samples were collected pre-exercise, after a 15 min warm-up, directly after peak exertion, and during the early recovery phase, enabling longitudinal assessment of rapid stress-induced molecular dynamics (Fig. 2A).

**Figure 2.**
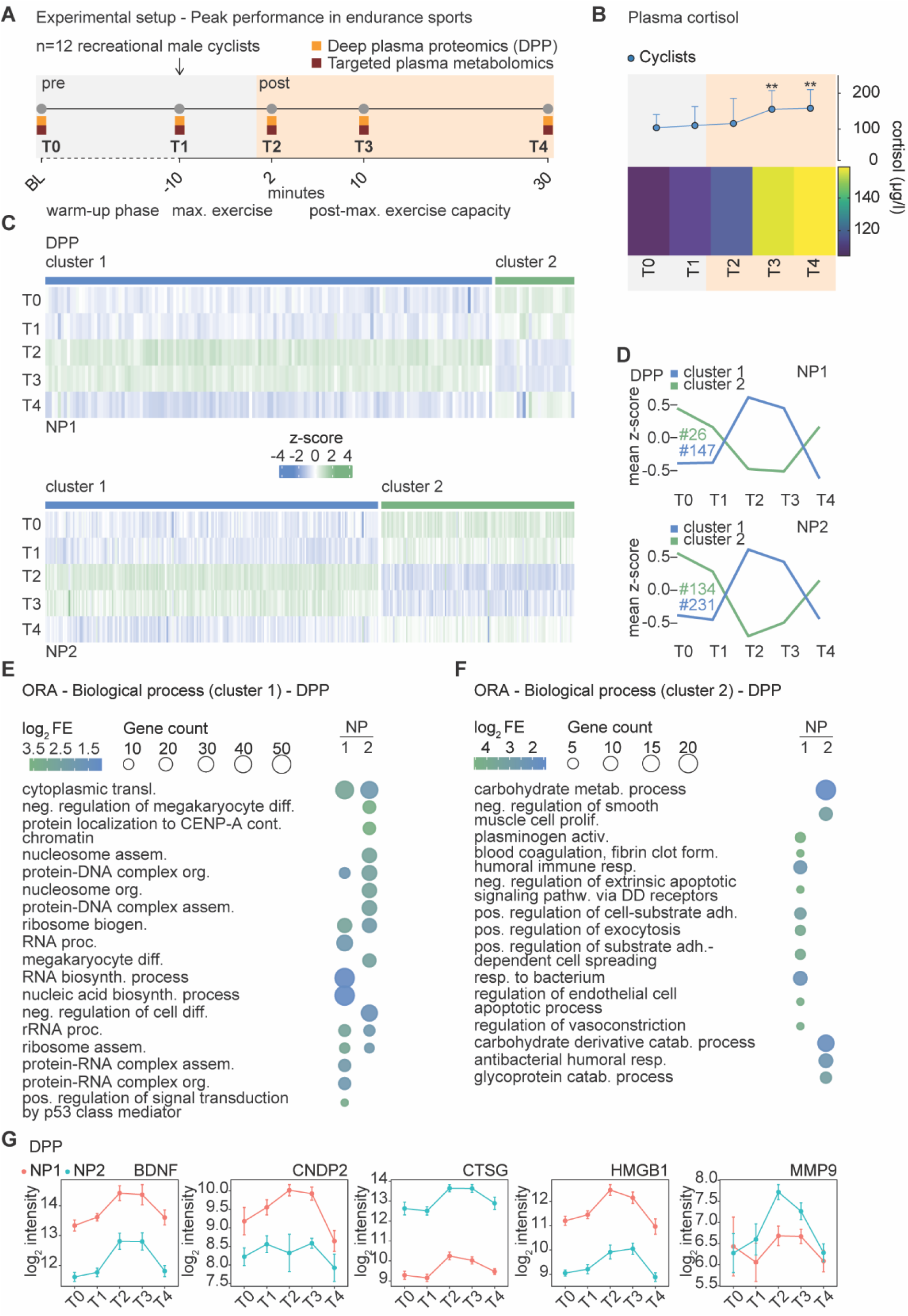
Physical stress induces coordinated plasma proteome remodeling. **(A)** Experimental design of the endurance sports cohort. n=15 recreationally active male participants performed a standardized incremental cycling test to volitional exhaustion (VO₂max). Plasma samples were collected at 5 timepoints (T0–T4): baseline (BL), after a 15-minute warm-up (T1), immediately after maximal exertion (T2), and during early recovery (T3, T4). Deep plasma proteomics (DPP) and targeted plasma metabolomics were performed at indicated timepoints. **(B)** Plasma cortisol levels (line plot, mean + SD) and heatmap of individual cortisol trajectories across timepoints. Shading indicates the post-exercise period. Linear mixed model with post-hoc comparisons of estimated marginal means against baseline (T0) using Dunnett-type contrasts; Holm–Bonferroni correction for multiple comparisons. n=8. * p < 0.05, ** p < 0.01, *** p < 0.001) **(C)** Heatmaps of z-scored protein abundances for significantly regulated proteins (n=12; adjusted p < 0.05; linear model with empirical Bayes moderation, blocking on subject; BH correction for multiple testing) across timepoints for NP1 (top) and NP2 (bottom). Proteins are grouped into two unsupervised clusters (cluster 1: blue; cluster 2: green). **(D)** Mean z-score trajectories for the two protein clusters across timepoints for NP1 (top) and NP2 (bottom). Numbers indicate the count of proteins per cluster. **(E)** Over-representation analysis (ORA) of biological process gene ontology terms for cluster 1. Results shown for NP1 and NP2. **(F)** ORA of biological process gene ontology terms for cluster 2. Results shown for NP1 and NP2. **(G)** Log₂ intensity trajectories (mean ± SEM) for selected stress-relevant proteins across timepoints for NP1 (red) and NP2 (teal).

Given the robust cortisol response observed during the TSST, we asked whether acute physical stress engages the HPA axis to a comparable or greater extent. To capture the hormonal landscape of the exercise response, we quantified plasma cortisol throughout the experiment (Fig. 2B). Exercise to exhaustion induced pronounced increases in cortisol, indicating robust activation of adrenal glucocorticoid pathways during acute physical stress.

To understand how these systemic hormonal signals translate into broader molecular adaptations, we next profiled the circulating plasma proteome following maximal exercise in 12 male participants. Deep proteomic profiling (Fig. S4A-F)) identified a large number of significantly regulated proteins across post-exercise timepoints compared to baseline (NP1: 101 proteins at T2 vs T0; NP2: 324 proteins at T2 vs T0; adjusted *p* < 0.05; Table S9, S10) (Fig. S4G). Among the most strongly regulated proteins were factors associated with immune activation and granule release (e.g. LTF, CTSG), stress-responsive alarmins (e.g. S100A8/A9), and cytoskeletal and intracellular stress regulators, reflecting rapid systemic adaptation to acute physical load.

To investigate how these exercise-responsive plasma proteins exhibited coordinated temporal dynamics, we performed unsupervised clustering of regulated proteins based on their abundance trajectories across the experiment. This analysis revealed two dominant and conserved response patterns (Fig. 2C, 2D). One cluster exhibited a rapid increase in protein abundance immediately after maximal exertion, followed by gradual return to baseline during recovery, whereas the second cluster showed reciprocal dynamics, characterized by transient decreases during peak stress and early recovery. The highly similar profiles observed across both NPs indicate that these patterns are robust and independent of nanoparticle chemistry. Gene ontology analysis of these opposing temporal clusters revealed complementary biological programs underlying the systemic stress response (Fig. 2E, 2F; Table S11). Proteins increased at peak exertion were enriched for RNA processing, ribosome biogenesis, and chromatin-associated processes, consistent with the release of intracellular biosynthetic and gene expression-related components during maximal physical load. In contrast, proteins showing reciprocal decreases were enriched for coagulation-, metabolic-, and humoral immune–related pathways, reflecting coordinated modulation of systemic plasma components during acute stress adaptation.

Within these coordinated response patterns, several proteins with established roles in stress and exercise physiology, exhibited pronounced and temporally structured regulation, including mediators of neurotrophic signaling (BDNF), metabolic regulation (CNDP2), innate immune activation (CTSG, HMGB1), and extracellular matrix remodeling (MMP9) (Fig. 2G).

Together, these results demonstrate that acute physical stress induces a rapid and highly organized remodeling of the circulating plasma proteome, characterized by conserved temporal dynamics and coordinated pathway-level regulation. Exercise-triggered increases of intracellular biosynthetic and gene expression-associated proteins occurred in parallel with transient reductions in coagulation-, metabolic-, and humoral immune-related components, indicating a coordinated shift from homeostatic maintenance toward acute stress adaptation. This pronounced systemic response contrasts sharply with the minimal plasma proteomic effects observed during psychological stress. These findings position controlled physical exercise as an intermediate stress modality, eliciting clear but largely transient plasma proteome remodeling compared with the minimal changes observed during psychological stress.

### Mixed psychological–physical stress induces amplified and sustained plasma proteome remodeling

Having defined systemic molecular responses to isolated psychological and physical stress, we next investigated a paradigm combining both modalities. We analyzed plasma samples from 22 male participants undergoing bungee jumping, a naturalistic paradigm combining acute psychological threat with intense physical load (Fig. 3A; Table S12). Blood samples were collected at baseline (7 days before the bungee jump), 50 and 15 minutes prior to the jump to capture anticipatory stress responses and at multiple timepoints following the jump (20, 60, 120 and 240 minutes), as well as at a late follow-up 7-10 days later. This longitudinal sampling scheme enabled assessment of both anticipatory and post-stress molecular dynamics across acute stress and recovery phases. Moreover, this setting enabled us to test whether simultaneous exposure to intense psychological threat and high mechanical load amplifies systemic molecular responses beyond those observed for either stress modality alone.

**Figure 3.**
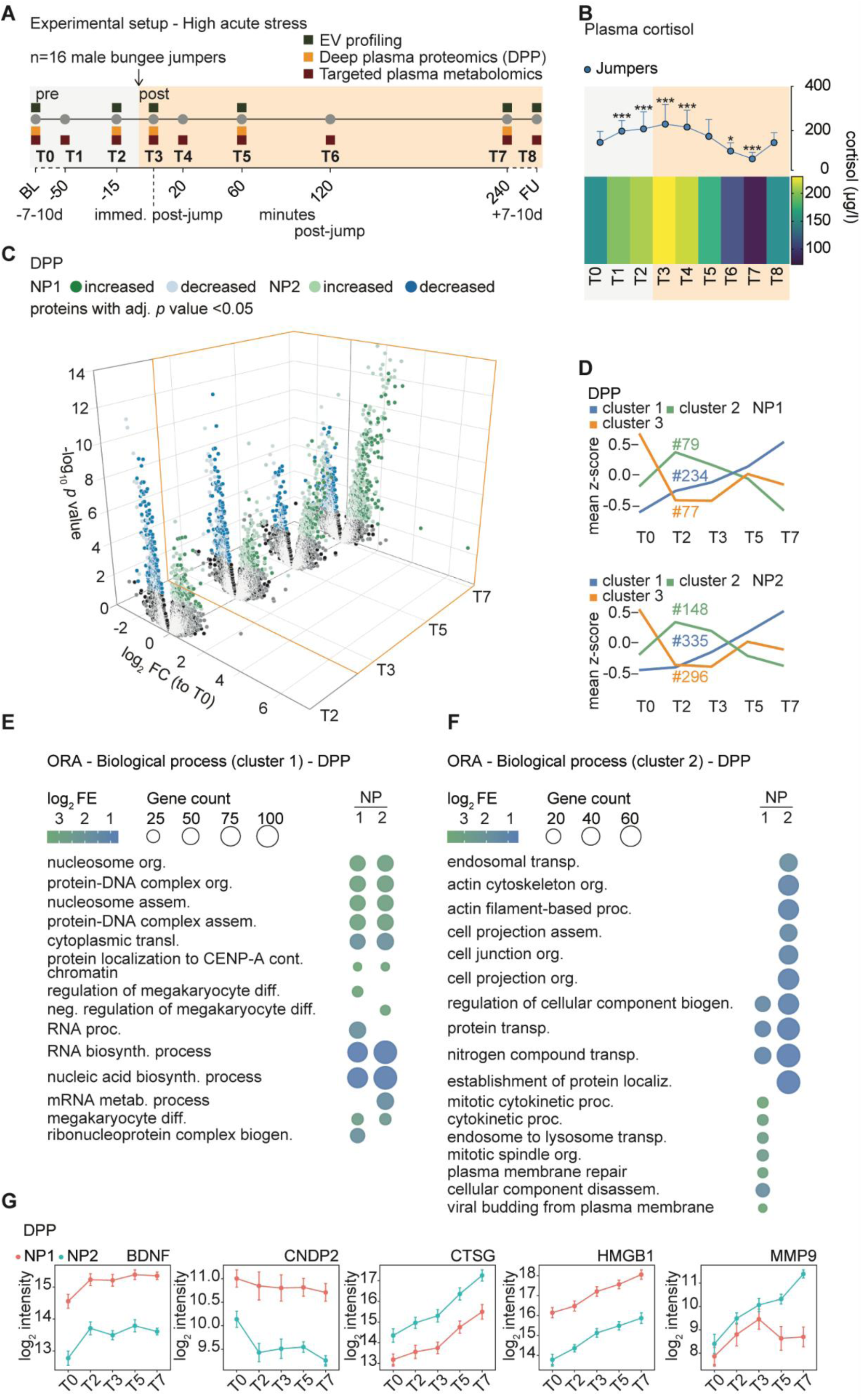
Mixed psychological-physical stress induces amplified and sustained plasma proteome remodeling. **(A)** Experimental design of the bungee jumping cohort. n=22 male bungee jumpers. Plasma samples were collected at 9 timepoints (T0–T8) spanning 7–10 days pre-jump (BL), anticipatory timepoints (T1, T2), immediately post-jump (T3), and up to 240 minutes post-jump (T4–T7), with a follow-up visit 7–10 days later (T8). EV profiling, deep plasma proteomics (DPP), and targeted plasma metabolomics were performed at indicated timepoints. **(B)** Plasma cortisol levels (line plot, mean + SD, n = 17-20 male bungee jumpers) and heatmap of individual cortisol trajectories across timepoints. Shading indicates the post-jump period. Linear mixed model with post-hoc comparisons of estimated marginal means against baseline (T0) using Dunnett-type contrasts; Holm–Bonferroni correction for multiple comparisons. * p < 0.05, ** p < 0.01, *** p < 0.001) **(C)** Three-dimensional volcano plot displaying differential protein abundance across post-jump timepoints (T2, T3, T5, T7) relative to baseline (T0) for NP1 and NP2. Significantly regulated proteins (n=16; adjusted p < 0.05; moderated paired t-test, BH correction) are color-coded by direction of regulation. **(D)** Mean z-score trajectories of significantly regulated proteins (n=16; adjusted p < 0.05; linear model with empirical Bayes moderation, blocking on subject; BH correction for multiple testing) for three unsupervised protein clusters across timepoints for NP1 (top) and NP2 (bottom). Numbers indicate protein counts per cluster. **(E)** ORA of biological process gene ontology terms for cluster 1. Results shown for NP1 and NP2. **(F)** ORA of biological process gene ontology terms for cluster 2. Results shown for NP1 and NP2. **(G)** Log₂ intensity trajectories (mean ± SEM) for selected stress-relevant proteins across timepoints for NP1 (red) and NP2 (teal).

Initially, we assessed whether our setup induced psychological stress in the participants. Self-report questionnaire responses indicated an increase in subjective stress, arousal, and enthusiasm among the jumpers, which declined by the end of the experiment (Fig. S5A). Critically, these subjective responses were mirrored at the endocrine level: plasma cortisol measurements confirmed a significant increase immediately after the jump, followed by a subsequent decline back toward baseline (Fig. 3B), providing converging evidence that bungee jumping reliably engaged both the psychological and physiological stress axes.

To determine whether the endocrine activation observed during bungee jumping translated into systemic proteomic changes, we next profiled the circulating plasma proteome across the experiment (Table S13). In contrast to the largely stable plasma proteome observed during purely psychological stress and the transient changes detected after maximal cycling effort controlled physical exercise, deep plasma proteomic profiling revealed extensive and highly coordinated remodeling of circulating proteins following bungee jumping, with widespread regulation that peaked 4 h post-jump (Fig. 3C; Table S14). Consistent with this escalating stress gradient, mixed psychological-physical stress produced the most extensive and long-lasting remodeling of the circulating proteome across all paradigms analyzed. Notably, many of the most strongly upregulated proteins, including HMGB1, MNDA and BPI, were nuclear factors with established extracellular functions in immune signaling pathways such as RAGE receptor binding.

To resolve the temporal organization of this response, we performed unsupervised clustering of significantly regulated proteins based on their abundance trajectories. This analysis uncovered three distinct and conserved response patterns that were highly similar across NPs (Fig. 3D; Table S15). The largest cluster (blue) displayed a continuous increase in protein abundance over time, reaching maximal regulation at T7, consistent with a dominant and sustained plasma proteomic response to mixed psychological–physical stress.

In addition to this delayed response, one cluster (green) showed maximal regulation already at the pre-jump timepoint (T2), indicating early proteomic changes preceding the physical stressor and coinciding with elevated cortisol levels. A third cluster (orange) exhibited reciprocal dynamics, characterized by a pronounced decrease at T2 and T3 followed by partial recovery, suggesting transient suppression of specific circulating proteins during the initial stress phase (Fig. 3D; Fig. S5H).

Gene ontology analysis of these clusters revealed a striking enrichment of intracellular biological processes (Fig. 3E, F; Fig. S5J; Table S16). Proteins in the dominant late-response cluster (blue) were strongly enriched for RNA biosynthesis, RNA processing, ribonucleoprotein complex assembly, and chromatin organization, indicating sustained release of nuclear proteins following mixed psychological–physical stress exposure (Fig. 3E). In contrast, the anticipatory cluster (green) was enriched for pathways related to protein localization, cytoskeletal and actin filament organization, endosomal transport, and plasma membrane–associated remodeling, consistent with early cellular reorganization and stress-induced secretory activity (Fig. 3F).

Several proteins with established roles in stress and exercise physiology exhibited pronounced and temporally structured regulation. Similar to the exercise paradigm, BDNF, CTSG, HMGB1, and MMP9 were strongly induced in response to this stressor, whereas CNDP2 was downregulated, in contrast to its upregulation during physical exercise (Fig. 3G). Notably, many of these upregulated proteins remained elevated even 4 hours after the jump, indicating a sustained systemic response, whereas the exercise-induced proteomic changes were largely transient (Fig. 3G). Consistent with this observation, the sustained release of MMP9 mirrors previous reports identifying this protein as a glucocorticoid-responsive secreted factor (Martinelli et al., 2021), suggesting convergent stress-induced secretory mechanisms across distinct experimental paradigms.

These findings demonstrate that mixed psychological-physical stress triggers a pronounced and sustained remodeling of the circulating plasma proteome. Compared with the minimal changes observed during purely psychological stress and the transient responses induced by controlled physical exercise, bungee jumping elicits a broader and longer-lasting (up to 4 hours) systemic proteomic response characterized by sustained release of nuclear stress signals and immune-associated proteins. Taken together, the three stress paradigms reveal a clear hierarchy of systemic proteomic responses: minimal changes during purely psychological stress, transient remodeling following physical exertion, and extensive, sustained proteome reorganization during combined psychological–physical stress.

### Shared and stressor-specific circulating metabolic and protein signatures across stress modalities

Having defined minimal proteomic responses to psychological stress, transient remodeling during maximal cycling effort,, and sustained proteome reorganization during mixed psychological–physical stress exposure, we next asked to what extent these responses share common molecular components across stress modalities.

To systematically compare molecular responses across stress modalities, we integrated plasma proteomic and metabolomic data obtained during psychological stress (TSST), physical exertion (bicycle ergometer test), and mixed psychological–physical stress (bungee jumping). This integrative analysis allowed us to determine which circulating molecular signatures are stressor-specific and which represent shared responses to acute stress exposure.

Consistent with the minimal proteomic remodeling observed during TSST, psychological stress resulted in only a small number of significantly regulated plasma proteins and showed little overlap with either physical or mixed stress paradigms, even when considering increasingly larger sets of top-ranked proteins (Fig. 4A). This limited overlap underscores that acute psychological stress primarily engages endocrine and metabolic signaling while leaving the circulating proteome largely stable. In contrast, physical exercise and bungee jumping displayed substantial convergence at the proteome level, sharing 205 significantly regulated proteins (Fig. 4B; Table S17). Peak-time comparisons further demonstrated that these shared proteins were regulated in the same direction across both paradigms, defining a coherent stress-associated core plasma proteome. Within the mixed psychological–physical stress paradigm, this core response was amplified in magnitude but largely conserved in composition and directionality relative to physical exercise (Fig. 4C; Fig. S6A). Thus, while exercise and bungee jumping differed in response amplitude and duration, they engaged largely overlapping systemic proteomic programs. Functionally, this shared stress-responsive core comprised intracellular stress sensors and alarmins (e.g. HMGB1/2/3, S100 family members), granule- and vesicle-associated immune effectors (ANXA1, BPI, CTSG), and proteins linked to cytoskeletal remodeling and unconventional secretion. These components point to coordinated activation of danger-associated signaling and secretory programs that propagate systemic stress responses through the circulation. Together, these analyses identify a conserved circulating stress-response proteome that is preferentially engaged by intense physiological stress and amplified during combined psychological–physical challenge, suggesting that systemic protein release represents a central molecular axis of human stress adaptation.

**Figure 4.**
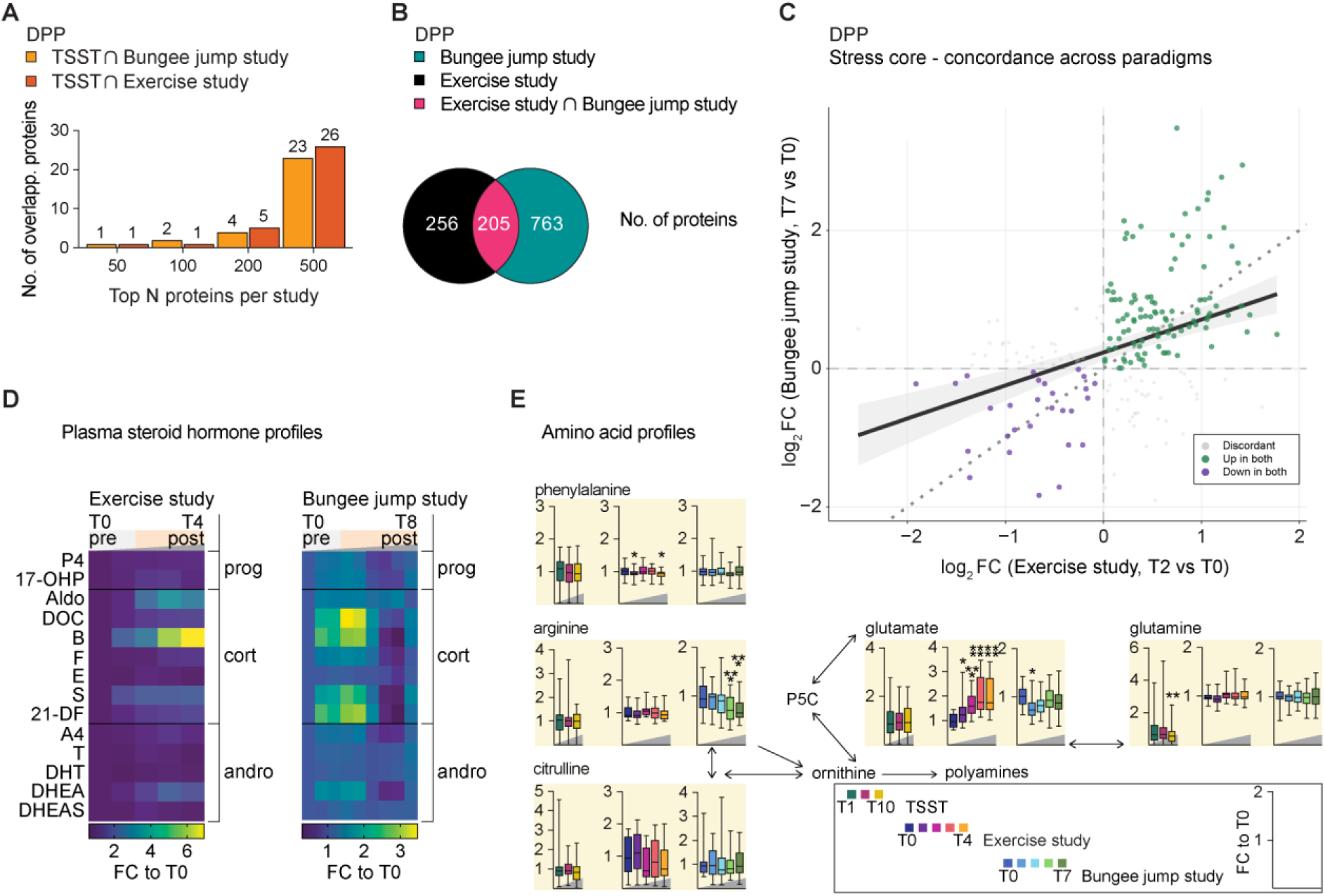
Shared and stressor-specific circulating proteomic and metabolic signatures across stress modalities. **(A)** Bar plot showing the number of overlapping top-ranked regulated proteins between the TSST and bungee jump study (orange) and between the TSST and exercise study (dark orange) at increasing protein set sizes (top 50, 100, 200, 500). Proteins were ranked by BH-adjusted p-value from time-course analysis (linear model with empirical Bayes moderation, blocking on subject; NP1 and NP2 collapsed by taking the minimum adjusted p-value per protein). **(B)** Venn diagram illustrating the overlap of significantly regulated proteins (BH-adjusted p < 0.05; time-course analysis by linear model with empirical Bayes moderation, blocking on subject) between the exercise study (black), bungee jump study (teal), and their intersection (=stress core). **(C)** Concordance scatter plot of NP-robust log₂ fold-change (minimum of NP1 and NP2) at fixed peak timepoints for all stress core proteins (n = 205; Exercise: T2 vs T0, Bungee jump: T7 vs T0). Each point represents one protein, colored by direction of regulation (green: upregulated in both; purple: downregulated in both; grey: discordant). The dotted line indicates the identity line (y = x); the solid line shows the linear regression fit (r = 0.38). Points above the identity line indicate greater fold-change in the bungee jump cohort. **(D)** Plasma steroid profiling heatmaps for the exercise cohort (left, T0–T4) and bungee jump cohort (right, T0–T8), showing fold-change to T0 for 14 steroids grouped by class: progestogens (prog), glucocorticoids (cort), and androgens (andro). Abbreviations: P4, progesterone; 17-OHP, 17-hydroxyprogesterone; Aldo, aldosterone; DOC, deoxycorticosterone; B, corticosterone; E, cortisone; S, 11-deoxycortisol; 21-DF, 21-deoxycortisol; A4, androstenedione; T, testosterone; DHT, dihydrotestosterone; DHEA, dehydroepiandrosterone; DHEAS, DHEA sulfate. Bungee study: n = 16-20; Sport study: n =7-8) **(E)** Targeted plasma metabolomics across stress paradigms. Metabolite changes in the arginine-citrulline-ornithine and glutamate/glutamine interconversion network. Box plots show fold-change to pre-stress timepoint across timepoints for TSST (FC to T1), endurance sports (FC to T0), and bungee jump (FC to T0) cohorts. Post-hoc comparisons against baseline: Bungee-Jumping and Exercise: estimated marginal means (emmeans) with Benjamini-Hochberg correction; TSST: Wilcoxon signed-rank test with Benjamini-Hochberg correction. (TSST: n=40; Exercise: n=15; Bungee: n=21-22; * p < 0.05, ** p < 0.01, *** p < 0.001; **** p < 0.0001)

To determine whether stressor-specific responses extend beyond the circulating proteome, we next examined metabolic adaptations across the different stress paradigms using plasma metabolomics (Table S18). Targeted analysis revealed coordinated remodeling of metabolic pathways that differed markedly in both breadth and magnitude across conditions. Physical stress paradigms induced the most extensive metabolic responses: in the exercise cohort, 24 of 35 measured neurotransmitter and amino acid metabolites were significantly altered (FDR < 0.05) (Fig. S7A), while in the bungee jump cohort, 12 of 47 metabolites reached significance (FDR < 0.05) (Fig. S7B,C). Psychosocial stress alone elicited a substantially more limited metabolic response, with only 2 of 56 measured metabolites significantly altered across all participants (FDR < 0.05) (Fig. S6A,D). Strikingly, this modest overall response masked a pronounced group difference: PD patients showed a markedly broader metabolic remodeling under psychological stress, with 6 metabolites reaching FDR-corrected significance, including N-lactoyl-phenylalanine (Lac-Phe) (Fig. S6C,F), whereas healthy controls showed virtually no significant metabolic changes (Fig. S6B,E). This pattern suggests that PD patients exhibit an exaggerated or dysregulated metabolic stress response to psychosocial challenge that is not observed in healthy individuals.

To illustrate the nature of these stress-induced metabolic changes, we examined the arginine–citrulline–ornithine axis and the glutamate/glutamine network, pathways well established as stress-responsive in the context of physiological challenge (Fig. 4E). These interconnected amino acid interconversion routes, linking arginine and citrulline via the urea cycle, ornithine to polyamine synthesis, and glutamate to glutamine through nitrogen buffering, showed consistent, time-dependent alterations under physical stress in both the exercise and bungee jump cohorts, while remaining largely unchanged under psychosocial stress alone (Fig. 4E).

Beyond metabolites, acute stress also triggers a rapid and coordinated steroidogenic response that varies in magnitude and duration across stressors. To capture the full breadth of the adrenal and gonadal steroidogenic response to physical and mixed stress, we performed targeted LC–MS/MS quantification of 14 circulating steroid hormones spanning three major classes, progestogens, glucocorticoids, and androgens, in the exercise and bungee jump cohorts (Fig. 4D). Both stress paradigms induced robust and class-specific steroidogenic responses. Glucocorticoids, particularly cortisol and corticosterone, showed the most pronounced increases, peaking immediately after maximal exertion (exercise cohort: T2) and shortly after the jump (bungee jump cohort: T3), consistent with canonical HPA axis activation. Progestogen intermediates, including 17-OHP and DOC, were co-regulated with cortisol, reflecting shared upstream steroidogenic flux through the adrenal cortex.

Androgen dynamics were more stressor specific. In the exercise cohort, DHEA and DHEAS showed transient increases following peak exertion, consistent with adrenal androgen co-secretion during sympathoadrenal activation, whereas testosterone and DHT remained relatively stable. In the bungee jump cohort, a more sustained androgen response was observed, with elevations persisting beyond the immediate post-stress period, potentially reflecting the prolonged adrenocortical activation characteristic of mixed psychological–physical stress.

### Mechanistic exploration of stress-induced secretion

To investigate the cellular and molecular mechanisms driving the pronounced and sustained plasma proteome remodeling observed after bungee jumping, we next focused on extracellular vesicles (EVs) as potential mediators of stress-induced protein secretion.

Semiquantitative profiling of EV subfractions using immunobead-based flow cytometry revealed a robust and sustained increase in several circulating EV populations following stress, peaking at 4 hours post-jump (T7). By the follow-up time point (T8, 7–10 days later), EV levels largely returned to baseline (T0), pointing towards a successful termination of stress-induced effects on EVs (Fig. 5A,B; Fig. S9A, B).

**Figure 5.**
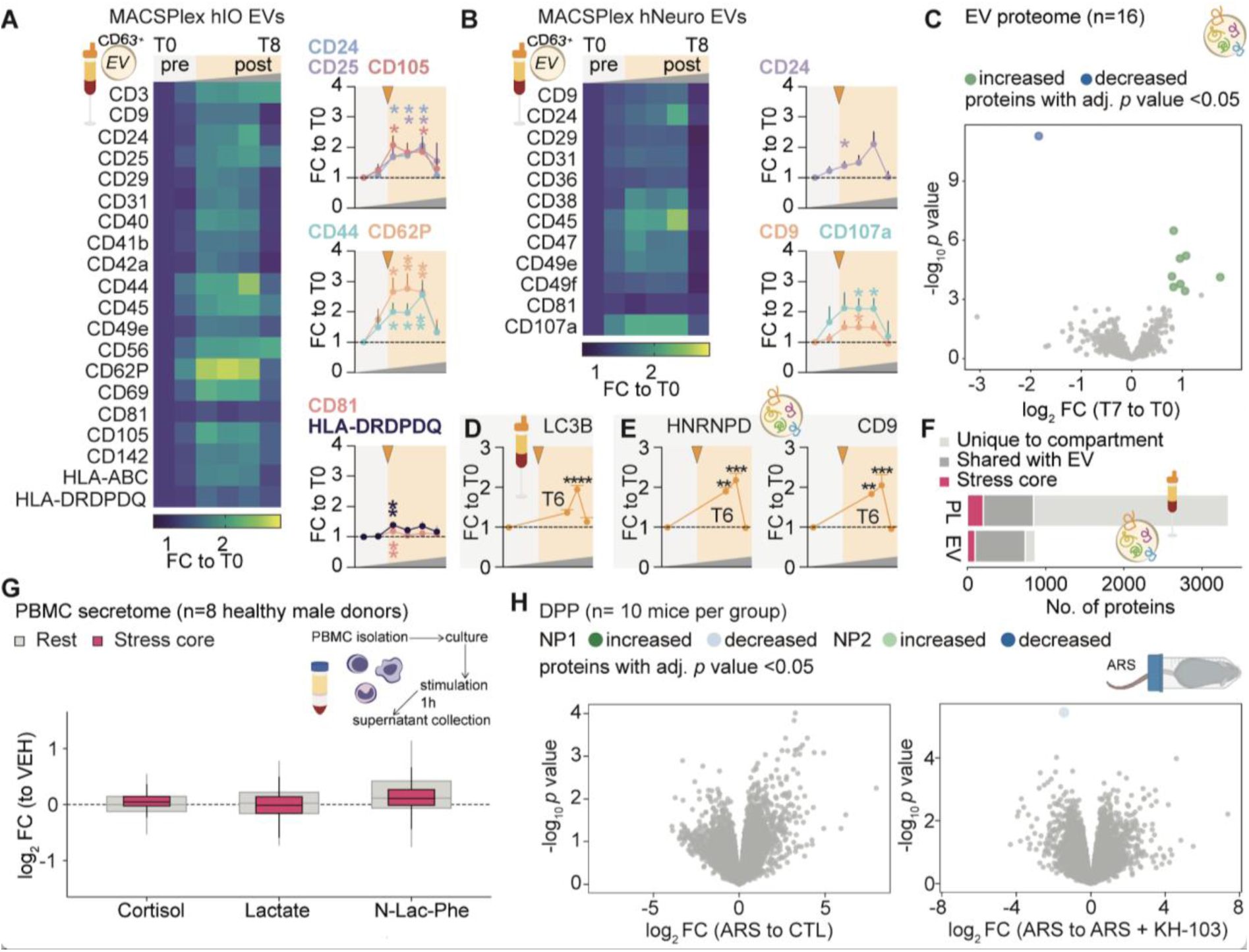
Stress-induced plasma proteome remodeling is driven by extracellular vesicle-mediated unconventional secretion. **(A)** Heatmap showing fold changes relative to baseline (T0) for different extracellular vesicle (EV) subpopulations in plasma of jumpers, identified by surface epitopes using the MACSPlex EV Kit for immune-oncology. Line plots display the mean of significantly regulated EV subpopulations over the course of the experiment (n = 14-18, Mixed effects model followed by Dunnett’s multiple comparisons test, *p* ≤ 0.05 (*), *p* ≤ 0.01 (**)). **(B)** Heatmap showing fold changes relative to baseline (T0) for EV subpopulations in plasma of jumpers identified by surface epitopes with the MACSPlex EV Kit for neurobiology. Line plots display the mean of significantly regulated EV subpopulations over the course of the experiment (n = 14-18, Mixed effects model followed by Dunnett’s multiple comparisons test *p* ≤ 0.05 (*). **(C** Volcano plot showing differential protein abundance in EV proteome of the bungee jumpers at T7 compared to baseline (T0) (n = 12; paired moderated t-tests; BH-adjusted p < 0.05). **(D)** Plasma LC3B levels (fold-change to T0, mean + SEM) across bungee jump timepoints, with the post-jump period shaded (n = 12, Friedman test followed by Dunn’s multiple comparisons, p ≤ 0.0001 (**\*\*\*\***)). **(E)** Targeted protein quantification of HNRNPD and CD9 in EV lysates (fold-change to T0, mean + SEM). (n = 12, Friedman test followed by Dunn’s multiple comparisons; p ≤ 0.01(**), p ≤ 0.001(***). **(F)** Bar plot comparing the number of proteins unique to the plasma compartment (PL), shared between plasma and EVs, or belonging to the stress core, for plasma (PL) and EV proteomes. **(G)** PBMC secretome experiment. Primary human PBMCs from n=8 healthy male donors were stimulated for 1 hour with cortisol, lactate, or N-lactoyl-phenylalanine (N-Lac-Phe), and supernatants were collected. Box plots show log₂ fold-change relative to vehicle (VEH) for stress core proteins (pink) and all other detected proteins (grey) for each stimulus condition. **(H)** Deep serum proteomics (DPP) in a mouse acute restraint stress (ARS) model (n=10 mice per group). Volcano plots show differential protein abundance for ARS vs. control (CTL, left) and ARS vs. ARS + KH-103 (glucocorticoid receptor degrader, right) using NP1 (green/light blue) and NP2 (dark green/dark blue). (Moderated t-tests; Benjamini-Hochberg (BH) correction for multiple testing; adjusted p < 0.05 highlighted).

Targeted protein analyses confirmed stress-associated increases in the canonical EV markers, including CD9 and the EV cargo protein HNRNPD in lysates of isolated EVs (Fig. 5 E). Additionally, plasma levels of LC3B, a central regulator of autophagy (Kabeya et al., 2000) and the loading of RNA-binding proteins into EVs (Leidal et al., 2020) were markedly elevated, providing a mechanistic link between autophagy-related pathways and stress-induced EV release (Fig. 5D).

Unbiased proteomic profiling of isolated EVs identified a distinct set of stress-regulated EV-associated proteins, with the largest changes observed at T7. Notably, these included immune-related factors such as CTSG (Fig. 5C). EV alterations were largely absent at other time points, mirroring the transient kinetics observed at the vesicle level (Fig. S9C, D; Table S19).

To place EV secretion into a systemic context, we compared EV proteomes and the previously defined stress-associated protein core. A substantial fraction of stress-regulated plasma proteins, including core factors detected in both exercise and bungee studies, were present in EVs, supporting EV-mediated unconventional secretion as a central driver of stress-induced plasma proteomic signatures (Fig. 5F; Table S20).

To begin dissecting the upstream signals responsible for stress-induced protein secretion, we asked whether individual circulating stress mediators are sufficient to recapitulate the observed stress response in vitro. Primary human PBMCs were stimulated with selected stress hormones and metabolites, including cortisol, lactate and Lac-Phe, which were strongly regulated in our experiments (Fig. 1C, 2B, 3B, 4D). These stimuli alone failed to recapitulate the robust secretory patterns observed in vivo, eliciting only weak and variable directional changes in stress-core and other proteins (Fig. 5G; Fig. S10A-C; Table S21). These findings argue against a simple hormonal trigger and suggest that the in vivo secretory response requires the convergence of multiple stress signals.

Finally, to assess whether human stress-associated blood signatures can be reproduced under controlled conditions and mechanistically dissected in vivo, we analyzed serum proteomes from mice subjected to acute restraint stress (ARS), an established model for inducing psychological stress (Schmidt, 2024). To directly test the role of glucocorticoid receptor (GR) signaling, a subset of animals was treated with a GR-PROTAC to induce targeted GR degradation prior to stress exposure (Gazorpak et al., 2023). Mirroring the limited observed response in purely psychological stress (TSST) in humans, ARS alone did not induce detectable serum proteomic remodeling, and GR degradation neither unmasked nor amplified stress-induced protein release (Fig. 5H; Table S22), indicating that neither psychological stress nor glucocorticoid signaling alone is sufficient to drive systemic secretion.

Together, these data support a model in which stress-induced unconventional secretion is most robustly triggered under conditions combining psychological and physical stress. Rather than being solely dependent on glucocorticoid signaling, systemic proteomic remodeling appears to arise from the integration of multiple stress signals, including metabolic, cellular, and mechanical cues, which converge on EV-mediated unconventional secretory pathways in a stressor-specific manner.

## Discussion

Adaptive responses to acute stress require the rapid integration of cognitive appraisal, endocrine signaling, metabolic adaptation, and immune activation. By combining deep plasma proteomics and targeted metabolomics across three complementary human stress paradigms, psychological stress (TSST), physical exertion, and mixed psychological-physical stress, our study reveals that the qualitative nature of a stressor critically determines whether systemic molecular remodeling occurs. These data define a gradient of circulating responses in which endocrine activation alone is insufficient to drive large-scale plasma proteome remodeling, whereas stressors imposing metabolic and mechanical load engage coordinated systemic secretory programs.

A striking observation is the dissociation between hormonal stress activation and circulating proteomic responses. Despite robust HPA axis activation, purely psychological stress induced by the TSST resulted in minimal plasma proteomic remodeling. This finding challenges the common assumption that glucocorticoid signaling alone orchestrates systemic stress responses and instead suggests that endocrine activation is dissociable from large-scale protein release into circulation, pointing to additional physiological inputs, such as metabolic or mechanical load, as critical drivers of systemic secretory responses. Notably, this compartmentalization was also evident in individuals with panic disorder, a population with heightened stress sensitivity (Abelson et al., 2007), but showed no amplified peripheral proteomic responses. This stability of the plasma proteome likely reflects strong systemic buffering of transient neuroendocrine stress signals. The few regulated proteins were predominantly intracellular stress- and cytoskeleton-associated factors, suggesting selective release from activated immune or vascular cells rather than widespread secretory responses. Together, these observations suggest psychological stress alone largely remains confined to neuroendocrine signaling, while systemic plasma remodeling requires more intense or traumatic experiences or additional physiological inputs. In contrast, both physical exercise and mixed psychological–physical stress triggered extensive plasma proteomic responses. Across these paradigms, we identified a conserved stress-associated protein core comprising intracellular stress sensors, immune granule components, and cytoskeletal regulators. Many proteins within this core, including HMGB family members, S100 proteins and neutrophil granule enzymes, belong to the class of leaderless proteins that lack classical signal peptides, suggesting secretion through unconventional pathways rather than canonical ER-Golgi route (Rammes et al., 1997; Gardella et al., 2002; Rabouille, 2017; Galluzzi et al., 2018). In addition, the enrichment of alarmins and neutrophil granule proteins raises the possibility that regulated cell-death-associated processes, such as pyroptosis, necroptosis or NETosis, contribute to their appearance in plasma. Many of these molecules function as damage-associated molecular patterns (DAMPs), mediating systemic danger signaling and immune activation during cellular stress and tissue strain.

Our integrated extracellular vesicle (EV) analyses provide convergent evidence supporting EV-mediated secretion as a major contributor to these circulating signatures. We observed a robust expansion of circulating EV subpopulations, substantial overlap between EV- and plasma-detected proteins, and enrichment of canonical EV cargo molecules in the bungee jump cohort. These observations are consistent with prior studies demonstrating that acute exercise stimulates EV release (Whitham et al., 2018; Brahmer et al., 2019). However, compared to EVs released during exercise, EVs released after bungee jump are more sustained within the circulation (over 4 h), while exercise EVs are cleared within 90 minutes (Brahmer et al., 2019).

The observed increase in plasma LC3B levels in the bungee jump cohort further suggests engagement of LC3-dependent vesicle loading and secretion (LDELS), an autophagy-related pathway that selectively incorporates RNA-binding and cytosolic proteins into EVs (Leidal et al., 2020). This mechanism provides a plausible explanation for the prominent enrichment of RNA-binding proteins and nuclear factors within the stress-regulated plasma proteome. Notably, several strongly regulated proteins in our dataset, including HMGB1 and MMP9, have previously been shown to undergo secretion through secretory autophagy pathways under inflammatory stress conditions (Martinelli et al., 2021; Hisaoka-Nakashima et al., 2024), providing mechanistic precedent for autophagy-dependent release of intracellular alarmins.

Beyond shared mechanisms, our data reveal pronounced temporal differences between stress modalities and thereby potentially diverging stress intensities. Physical exercise induced rapid plasma proteomic changes that largely resolved during the recovery phase, consistent with the transient metabolic demands of acute exertion. In contrast, mixed psychological–physical stress elicited responses of greater amplitude and persistence, with several proteins remaining elevated for several hours after exposure. The presence of anticipatory molecular changes prior to the jump demonstrate that perceived threat alone can initiate early systemic reorganization before the physical challenge occurs. Such anticipatory responses have been described at the endocrine and neural level but have not been observed in circulating molecular profiles (Engert et al., 2013; Herman et al., 2016). This sustained elevation of alarmins and immune-associated proteins may have important physiological implications for long-term health. Prolonged circulation of DAMP molecules such as HMGB proteins and S100 family members has been implicated in neuroimmune communication and systemic inflammatory priming (Andersson et al., 2018; Yang et al., 2020). Such mechanisms could contribute to the link between repeated stress exposure and long-term vulnerability to cardiovascular, metabolic, and psychiatric disorders (Vaccarino & Bremner, 2024; Kivimäki et al., 2023; Slavich & Irwin, 2014).

Targeted metabolomics provided independent support for the stressor-specific molecular hierarchy observed at the proteome level. Physical stress paradigms induced broad and coordinated metabolic remodeling, with many measured metabolites significantly altered in both the exercise and bungee jump cohorts, whereas psychological stress alone elicited minimal metabolic changes in healthy individuals. This convergence between proteomic and metabolic responses reinforces the notion that systemic molecular remodeling requires physiological inputs beyond endocrine activation alone. Notably, the metabolic response to psychological stress differed markedly between healthy controls and individuals with panic disorder: while controls showed virtually no significant metabolic changes under the TSST, PD patients exhibited a substantially broader metabolic response, with significant alterations in both glycolytic and amino acid metabolites. This dissociation, amplified metabolic reactivity in the absence of differences in either cortisol or proteomic responses, suggests that panic disorder is associated with altered peripheral metabolic stress sensitivity through mechanisms that remain to be determined.

Our mechanistic perturbation experiments indicate that no single mediator is sufficient to reproduce the observed secretory response. Neither stimulation of immune cells with cortisol, lactate, or Lac-Phe, nor acute restraint stress in mice, even in the context of pharmacological glucocorticoid receptor degradation, recapitulate the extensive plasma remodeling observed in humans during physical or mixed stress. These findings suggest that systemic stress secretion requires integration of multiple converging physiological signals, including mechanical strain, metabolic flux, and endocrine signaling, which are most strongly engaged under conditions combining psychological and physical challenge.

Several limitations warrant consideration. First, the absence of a crossover design precluded within-individual comparisons across stress paradigms, which would allow more precise dissection of stressor-specific molecular effects. Second, the cellular origins of stress-regulated plasma proteins remain incompletely resolved. Although EV signatures suggest contributions from immune and potentially muscle-derived compartments, future studies integrating spatial transcriptomics or tissue-specific EV tracing will be required to further define the precise sources of systemic stress signals. Third, both sexes were represented in the TSST experiment, whereas the other two paradigms included only male participants, which may influence the generalizability of the findings.

In summary, our results demonstrate that acute stress is encoded in the circulating molecular landscape in a modality-dependent manner. While psychological stress remains largely confined to neuroendocrine signaling, physical and mixed stressors activate coordinated systemic programs characterized by EV-associated unconventional protein secretion and the release of a conserved stress-associated protein core. These findings establish a framework for understanding how diverse environmental challenges shape our circulating molecular landscape and highlight extracellular vesicle-mediated secretion as a central mechanism linking cellular stress states to systemic communication.

---

**Figure S1.**
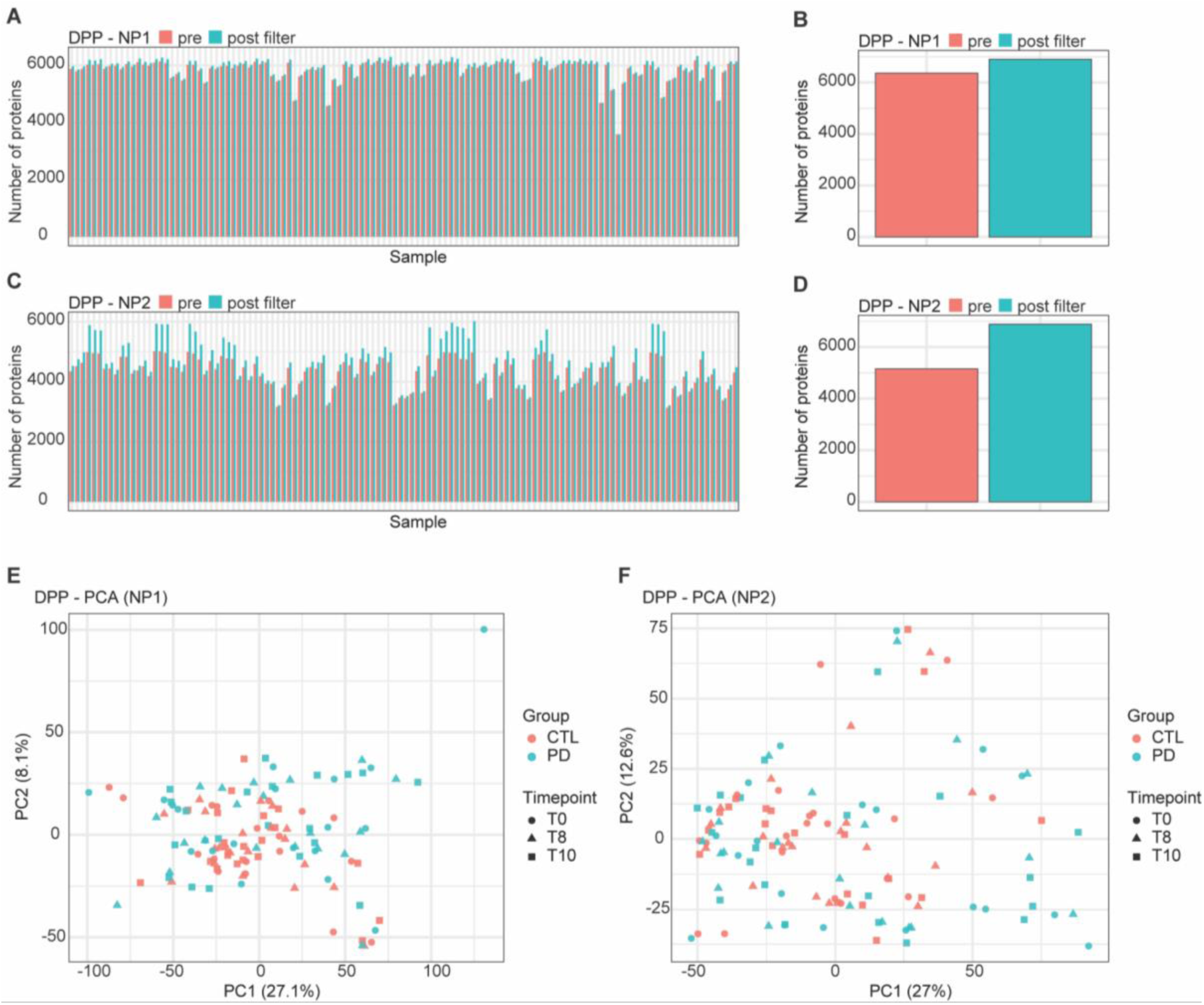
Deep plasma proteomics workflow characterization in the TSST cohort (n=40). **(A)** Number of measured proteins per sample in NP1 pre- and post-filtering. **(B)** Box plot showing the mean of measured proteins in NP1 pre- and post-filtering. **(C)** Number of measured proteins per sample in NP2 pre- and post-filtering. **(D)** Box plot showing the mean number of measured proteins in NP2 pre- and post-filtering. **(E)** Principal component analysis of NP1 measurements. **(F)** Principal component analysis of NP2 measurements.

**Figure S2.**
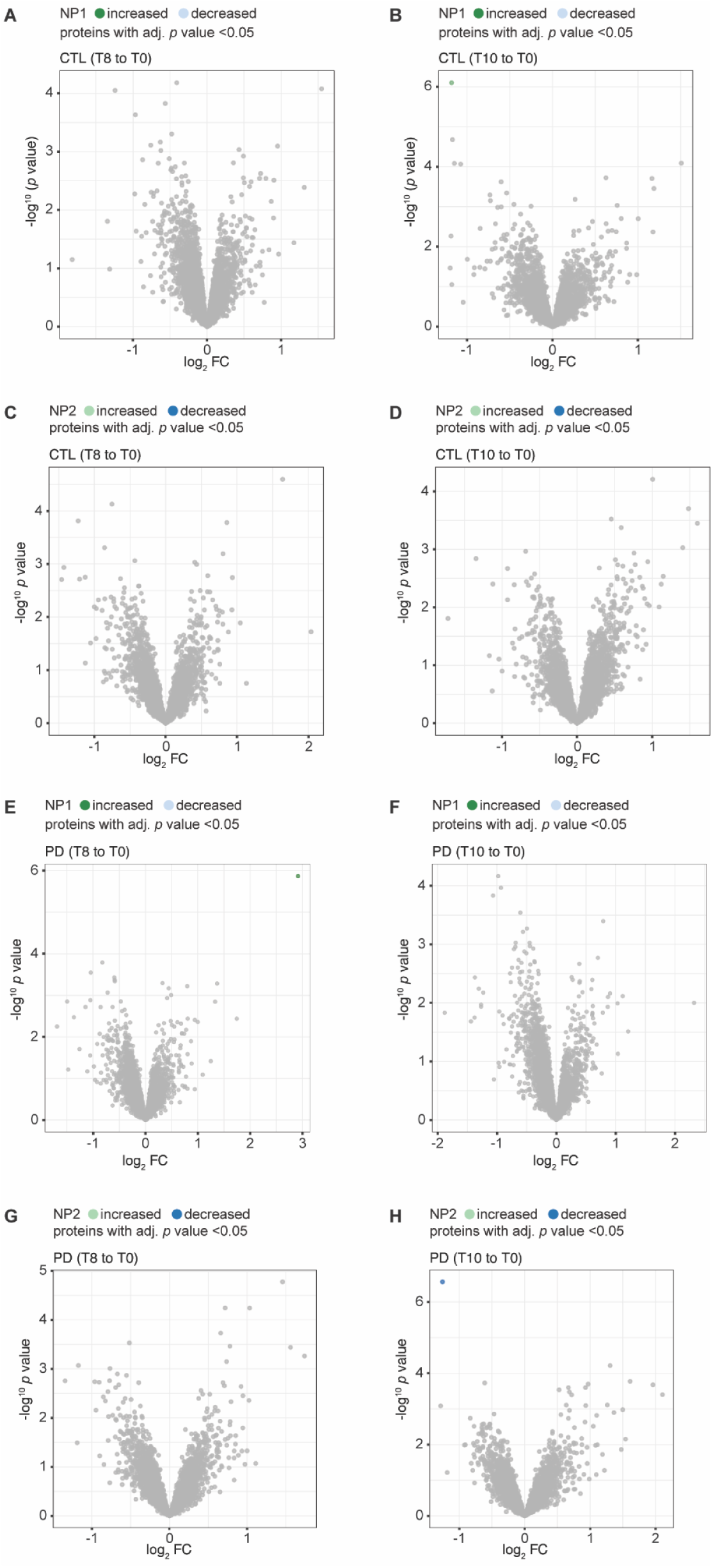
Differential plasma protein abundance across individual TSST timepoints in healthy controls and individuals with panic disorder. (A–H) Volcano plots showing differential protein abundance at individual post-TSST timepoints relative to pre-stress baseline (T0), analyzed separately for healthy controls (CTL; n=20) and individuals with panic disorder (PD; n=20) using NP1 and NP2 nanoparticle fractions. Moderated paired t-tests with Benjamini–Hochberg (BH) correction for multiple testing; significantly regulated proteins (adjusted p < 0.05) are highlighted.

**Figure S3.**
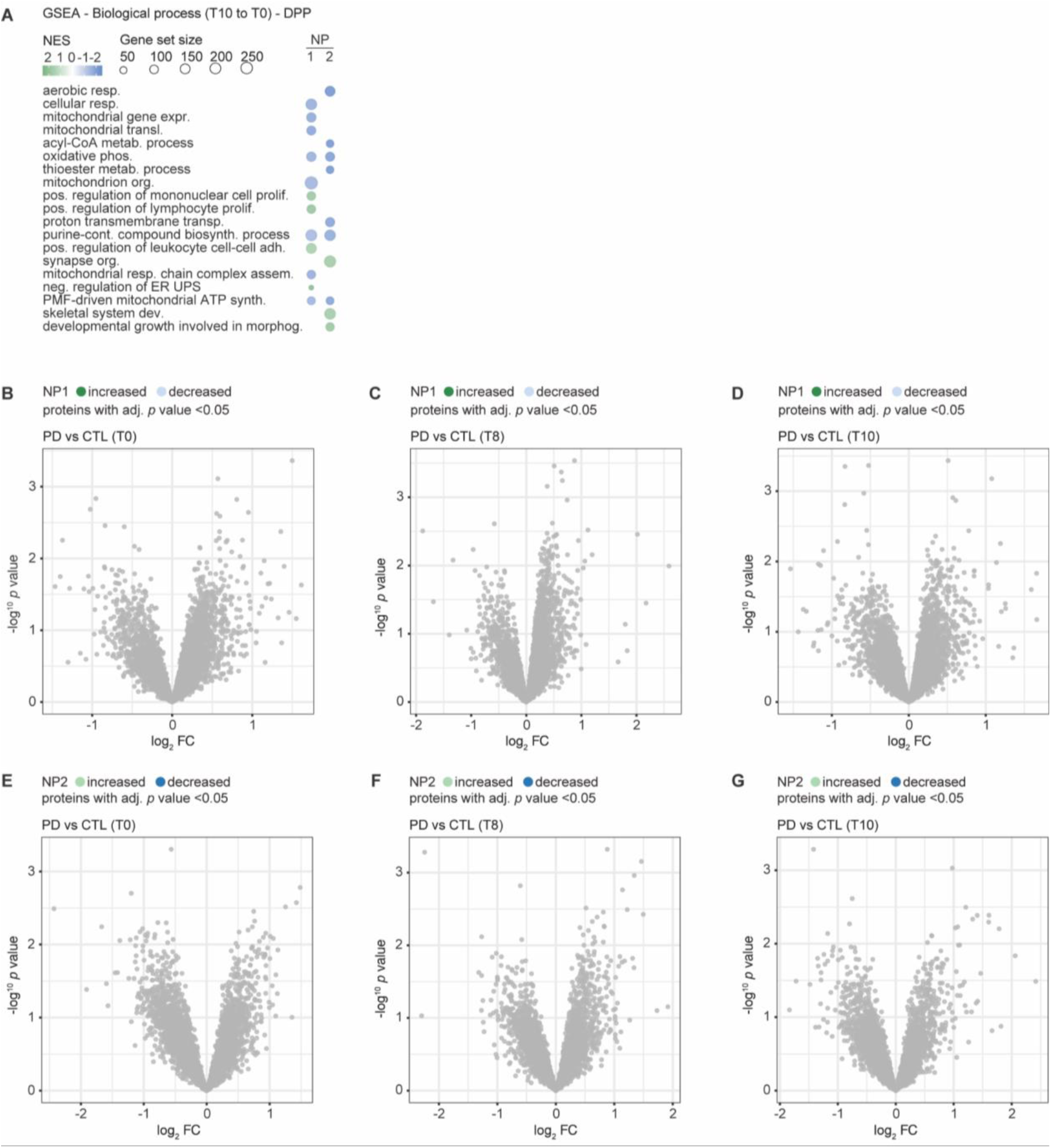
Gene set enrichment analysis and group comparison of plasma proteomic responses to psychological stress. (A) Gene Set Enrichment Analysis (GSEA) of ranked proteomic data (T10 vs. T0) for biological process gene sets, performed on the combined cohort. (B–G) Differential plasma protein abundance comparing individuals with panic disorder (PD) and healthy controls (CTL) at individual TSST timepoints. Moderated t-tests with BH correction; adjusted p < 0.05 highlighted.

**Figure S4.**
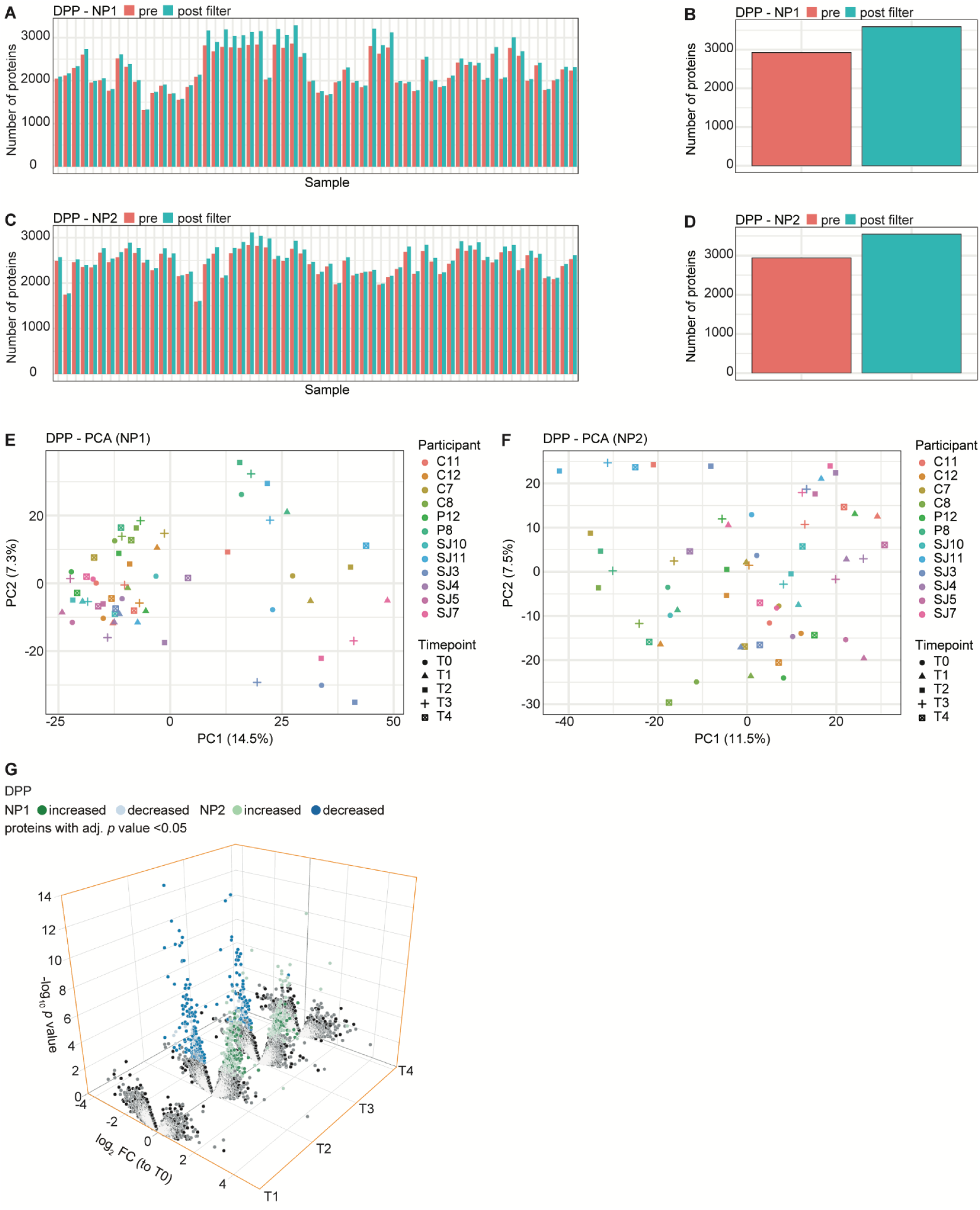
Deep plasma proteomics workflow characterization in the exercise cohort (n=12). **(A)** Number of measured proteins per sample in NP1 pre- and post-filtering. **(B)** Box plot showing the mean of measured proteins in NP1 pre- and post-filtering. **(C)** Number of measured proteins per sample in NP2 pre- and post-filtering. **(D)** Box plot showing the mean number of measured proteins in NP2 pre- and post-filtering. **(E)** Principal component analysis of NP1 measurements. **(F)** Principal component analysis of NP2 measurements. **(G)** Three-dimensional volcano plot displaying differential protein abundance across post-baseline timepoints (T1, T2, T3, T4) relative to baseline (T0) for NP1 and NP2. Significantly regulated proteins (n=12; adjusted p < 0.05; moderated paired t-test, BH correction) are color-coded by direction of regulation.

**Figure S5.**
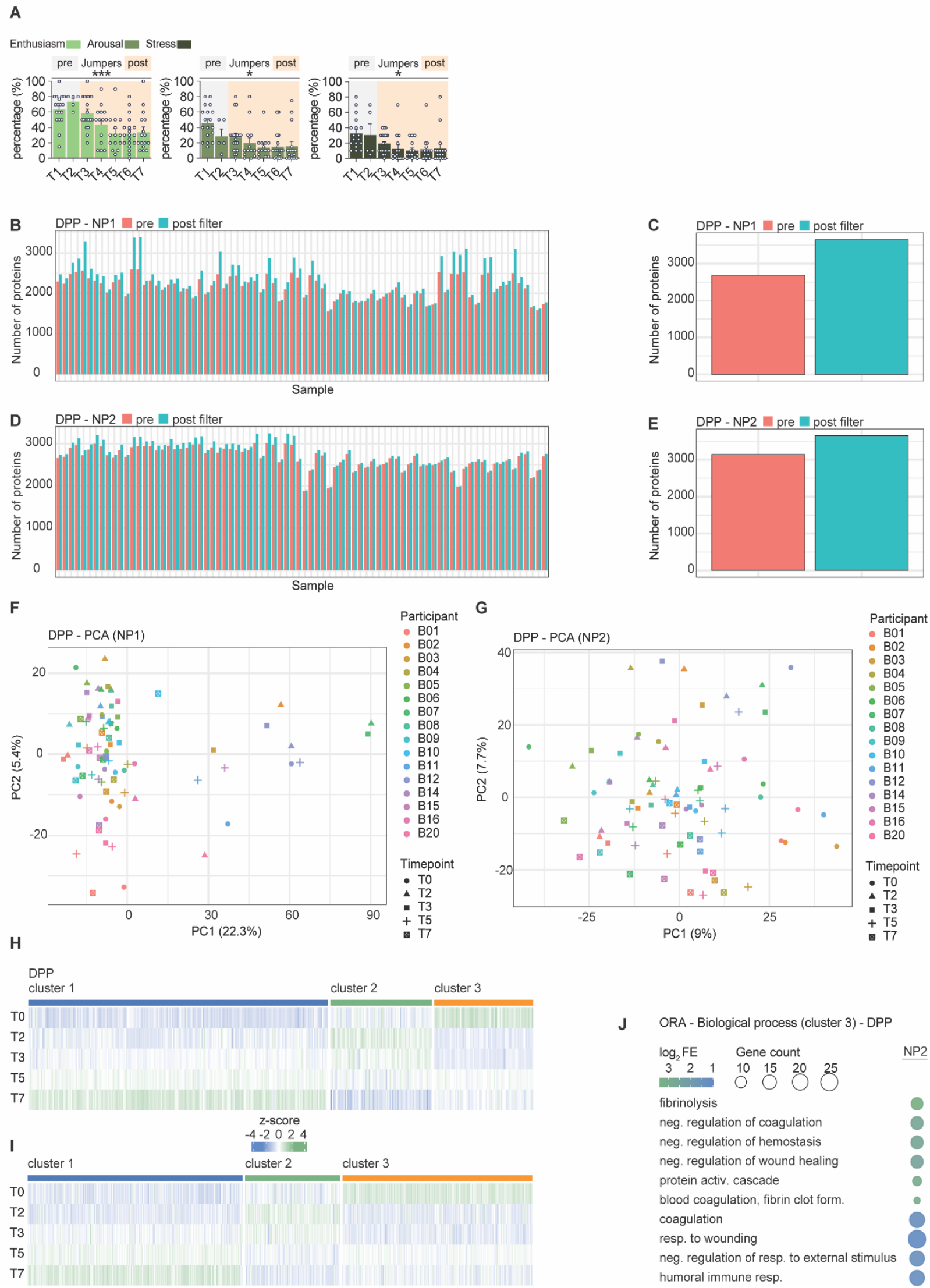
Deep plasma proteomics workflow characterization in the bungee jump cohort. **(A)** Subjective measures of enthusiasm, arousal and stress among jumpers on the day of the jump, as assessed by the “Single Stress Items” questionnaire (Mixed effects model, *p* ≤ 0.05 (*), *p* ≤ 0.001 (***)). **(B)** Number of measured proteins per sample in NP1 pre- and post-filtering. **(C)** Box plot showing the mean of measured proteins in NP1 pre- and post filtering. **(D)** Number of measured proteins per sample in NP2 pre- and post filtering. **(E)** Box plot showing the mean number of measured proteins in NP2 pre- and post-filtering. **(F)** Principal component analysis of NP1 measurements. **(G)** Principal component analysis of NP2 measurements. **(H,I)** Heatmap of z-scored protein abundances for significantly regulated proteins (n=16; adjusted p < 0.05; linear model with empirical Bayes moderation, blocking on subject; BH correction for multiple testing) across timepoints for **(H)** NP1 and **(I)** NP2. **(J)** ORA of biological process gene ontology terms for cluster 3. Results shown for NP2.

**Figure S6.**
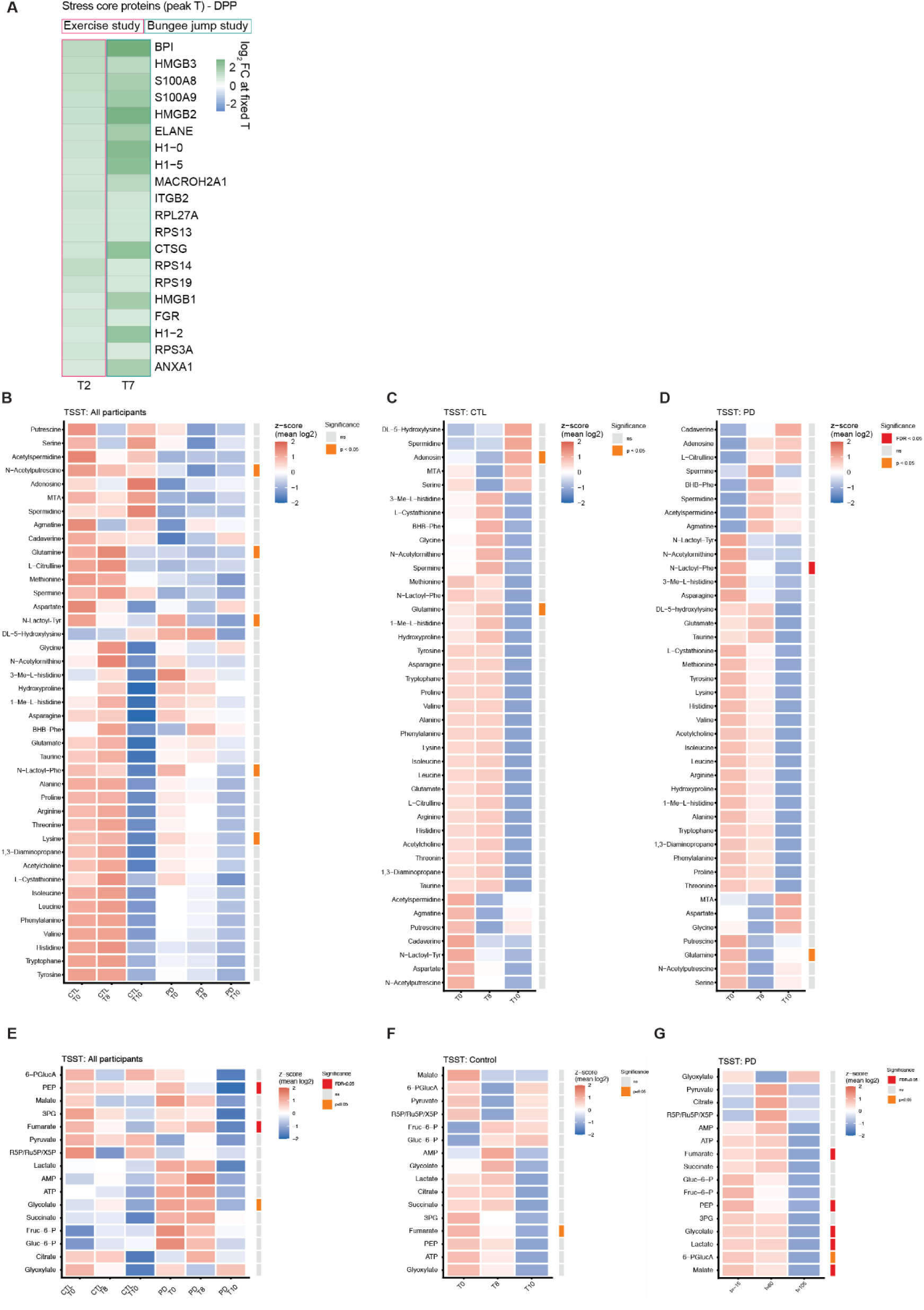
Targeted plasma metabolomics in the TSST cohort. **(A)** Heatmap of log₂ fold-change at peak timepoints (Sport: T2, Bungee: T7) for the top 20 proteins of the stress core (proteins significantly regulated in both cohorts, BH-adjusted p < 0.05). Effect sizes are NP-robust (minimum log₂ fold-change across NP1 and NP2). Proteins are ranked by the minimum NP-robust log₂ fold-change across Sport and Bungee. **(B)** Heatmap of log₂-transformed metabolite intensities across both groups in the TSST cohort, z-scored per metabolite (n=40; Friedman test, BH correction for multiple testing, custom neurotransmitter and amino acid panel (NT/AA)). The significance strip on the right indicates the result of the overall time effect test (red = FDR < 0.05, orange = p < 0.05 nominal, grey = p > 0.05) **(C)** Heatmap of log₂-transformed metabolite intensities (custom NT/AA panel) across the control group in the TSST cohort, z-scored per metabolite (n=20; Friedman test, BH correction for multiple testing). **(D)** Heatmap of log₂-transformed metabolite intensities (custom NT/AA panel) across the PD group in the TSST cohort, z-scored per metabolite (n=20; Friedman test, BH correction for multiple testing). **(E)** Heatmap of log₂-transformed metabolite intensities across both groups in the TSST cohort, z-scored per metabolite (n=40; Friedman test, BH correction for multiple testing, custom sugar phosphates and organic acid panel (SuP/OA)). The significance strip on the right indicates the result of the overall time effect test (red = FDR < 0.05, orange = p < 0.05 nominal, grey = p > 0.05) **(F)** Heatmap of log₂-transformed metabolite intensities (custom sugar phosphates and organic acid panel (SuP/OA)) across the control group in the TSST cohort, z-scored per metabolite (n=20; Friedman test, BH correction for multiple testing). **(G)** Heatmap of log₂-transformed metabolite intensities (custom sugar phosphates and organic acid panel (SuP/OA)) across the PD group in the TSST cohort, z-scored per metabolite (n=20; Friedman test, BH correction for multiple testing).

**Figure S7.**
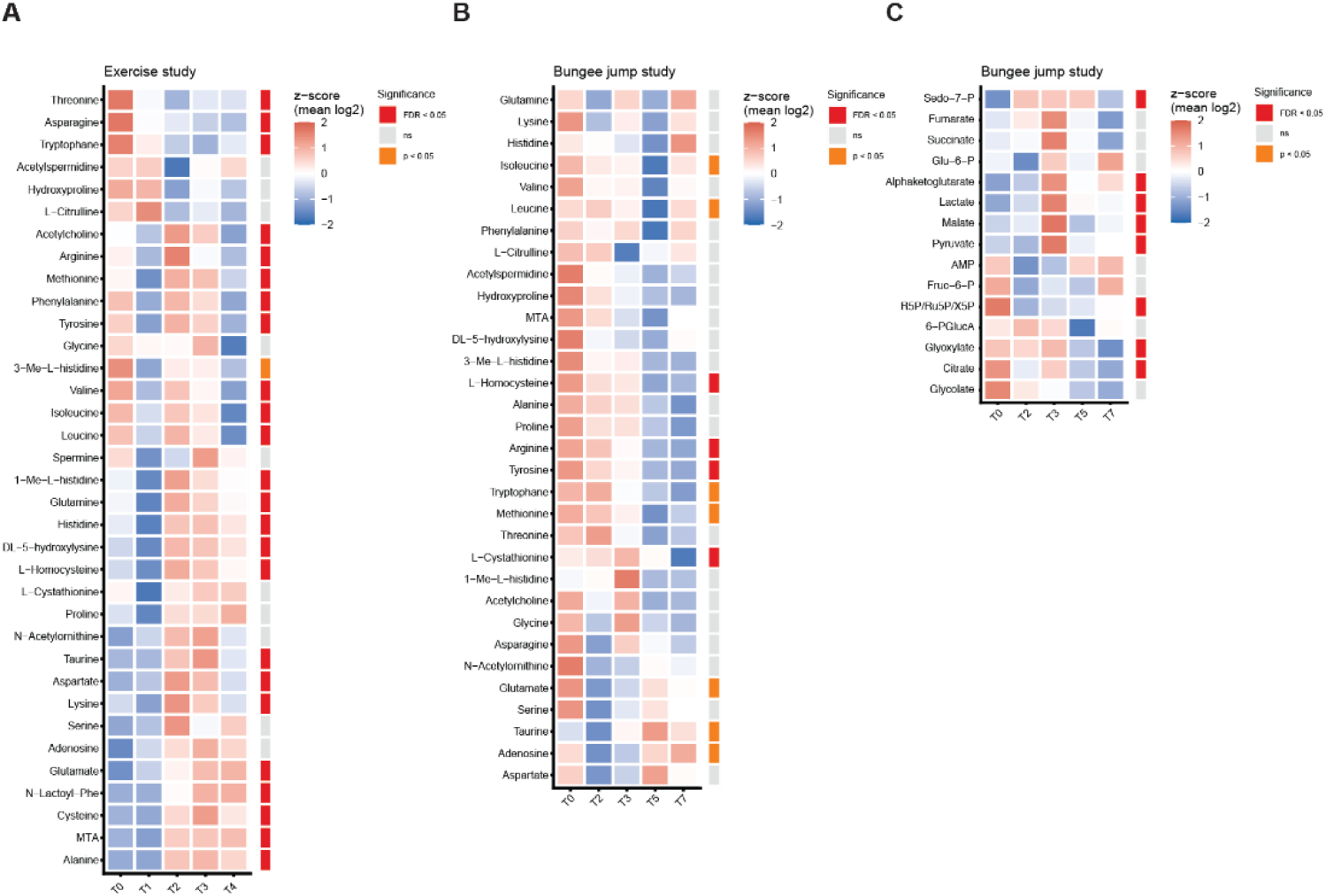
Targeted plasma metabolomics in the exercise and bungee jump cohort. **(A)** Heatmap of log₂-transformed metabolite intensities (custom neurotransmitter and amino acid panel (NT/AA)) in the exercise cohort, z-scored per metabolite (n=15; linear mixed model, BH correction for multiple testing). The significance strip on the right indicates the result of the overall time effect test (red = FDR < 0.05, orange = p < 0.05 nominal, grey = p > 0.05) **(B)** Heatmap of log₂-transformed metabolite intensities (custom NT/AA panel) in the bungee jump cohort, z-scored per metabolite (n=21-22; linear mixed model, BH correction for multiple testing). **(C)** Heatmap of log₂-transformed metabolite intensities (custom SuP/OA panel) in the bungee jump cohort, z-scored per metabolite (n=21-22; linear mixed model, BH correction for multiple testing). The significance strip on the right indicates the result of the overall time effect test (red = FDR < 0.05, orange = p < 0.05 nominal, grey = ns).

**Figure S8.**
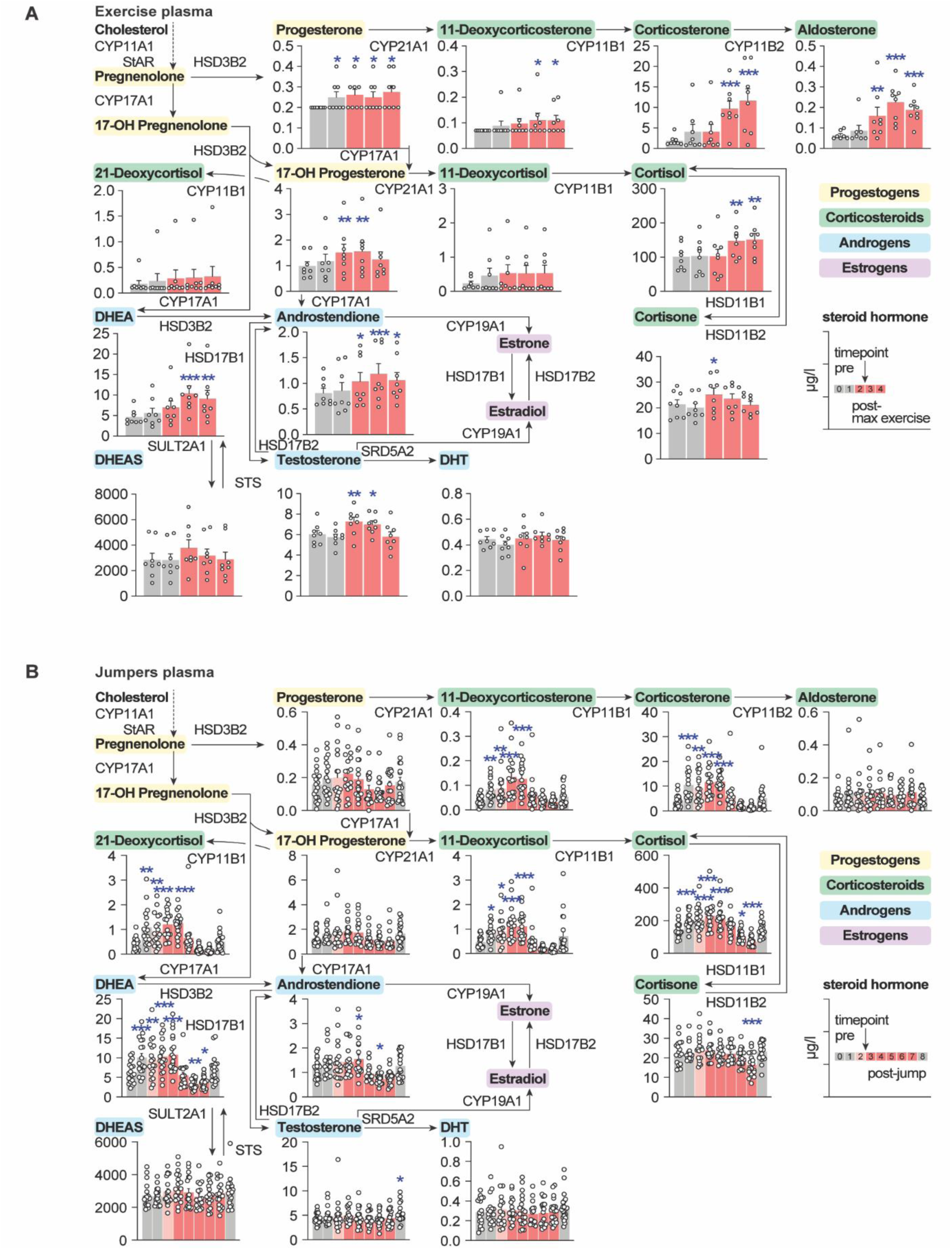
**(A)** Bar graphs showing concentrations of steroid hormones in the plasma at different timepoints within the exercise cohort (n = 7-8, Linear mixed models (random intercept per subject) with BH FDR correction across all 14 steroids. Post-hoc Dunnett-type contrasts vs. baseline (T0), Holm–Bonferroni corrected. * p < 0.05, ** p < 0.01, *** p < 0.001). (**B)** Bar graphs showing concentrations of steroid hormones in the plasma at different timepoints within the jumpers (n = 16-20, Linear mixed models (random intercept per subject) with BH FDR correction across all 14 steroids. Post-hoc Dunnett-type contrasts vs. baseline (T0), Holm–Bonferroni corrected. * p < 0.05, ** p < 0.01, *** p < 0.001).

**Figure S9.**
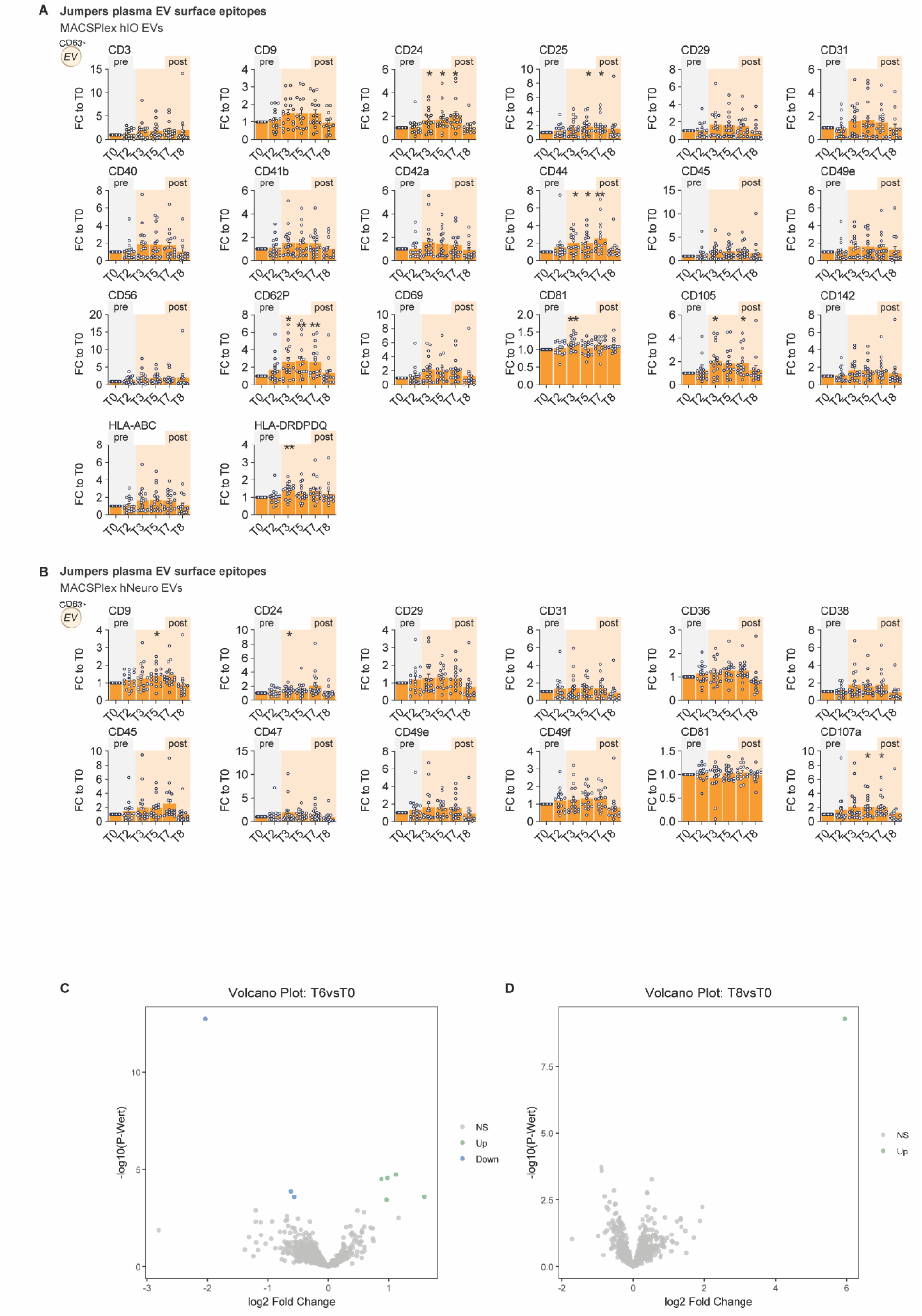
(A) Bar plots showing fold changes relative to baseline (T0) for different EV subpopulations identified by surface epitopes with the MACSPlex EV Kit for immune-oncology. (n = 14-18, Mixed effects model followed by Dunnett’s multiple comparisons test. (*p* ≤ 0.05 (*), *p* ≤ 0.01 (**), *p* ≤ 0.0001 (****)). **(B)** Bar plots showing fold changes relative to baseline (T0) for different EV subpopulations identified by surface epitopes with the MACSPlex EV Kit for neurobiology. (n = 14-18, Mixed effects model followed by Dunnett’s multiple comparisons test. (*p* ≤ 0.05 (*), *p* ≤ 0.01 (**), *p* ≤ 0.001 (***)). **(C)** Volcano plot showing differential protein abundance in EV proteome of the bungee jumpers at T6 compared to baseline (T0). (n = 12; paired moderated t-tests; BH-adjusted p < 0.05). **(D)** Volcano plot showing differential protein abundance in EV proteome of the bungee jumpers at T8 compared to baseline (T0). (n = 12; paired moderated t-tests; BH-adjusted p < 0.05).

**Figure S10.**
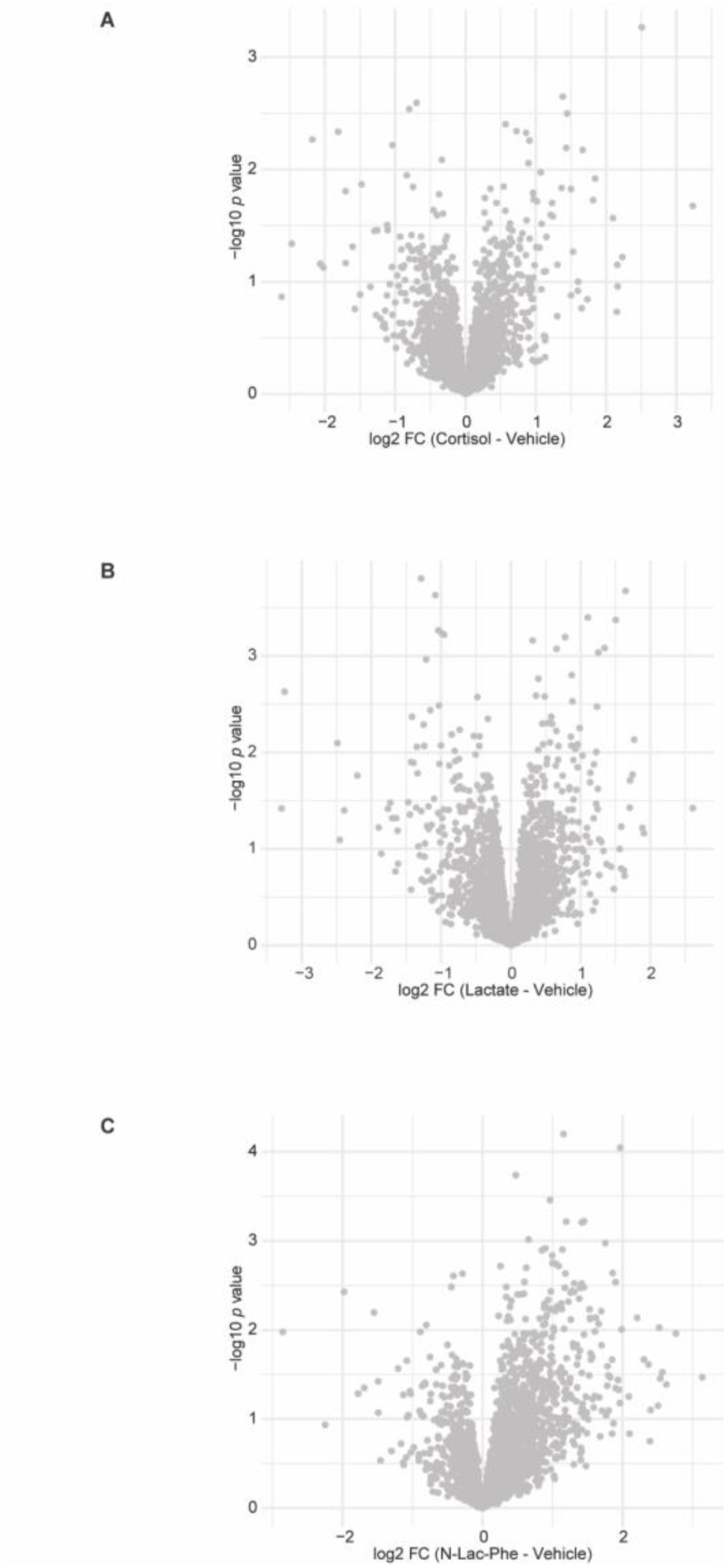
PBMC secretome responses to individual stress mediators. **(A)** Volcano plot showing differential protein abundance in the secretome of cells stimulated with cortisol compared to vehicle. (n = 8; moderated t-tests; BH-adjusted p < 0.05). **(B)** Volcano plot showing differential protein abundance in the secretome of cells stimulated with lactate compared to vehicle. (n = 8; moderated t-tests; BH-adjusted p < 0.05). **(C)** Volcano plot showing differential protein abundance in the secretome of cells stimulated with Lac-Phe compared to vehicle. (n = 8; moderated t-tests; BH-adjusted p < 0.05).

## Material and Methods

### Human studies

#### Cohort 1: Trier Social Stress Test

##### Ethical approval and study participants

Participants were recruited through newspaper advertisements, online announcements, and notice boards at various universities. Prior to enrollment, eligibility was assessed via a structured interview based on the Structured Clinical Interview for Axes I and II for DSM-IV (Wittchen et al., 1997; APA, 2000);. The experimental group consisted of individuals diagnosed with panic disorder according to DSM-IV criteria, aged between 18 and 65 years. A matched control group of healthy individuals was established, paired by age and gender to the experimental group. Groups did not differ significantly in age, gender, or use of oral contraceptives (all p > 0.05). Exclusion criteria included any acute or chronic comorbid medical illness, additional psychiatric diagnoses, current psychotropic medication or substance use, known allergies, and stressful life events within the preceding 6 months. Each group comprised n = 32 participants. All participants provided written informed consent. The study protocol was approved by the Ethics Committee of the Landesärztekammer Rheinland-Pfalz, Germany (No#2019-14188).

##### General study design

We conducted a single-center, non-blinded, non-randomized, controlled repeated-measures prospective study to investigate molecular and psychological responses to acute psychosocial stress in healthy controls and in panic disorder patients. All assessments were performed on weekdays between 2:00 and 6:00 p.m. The participants were instructed not to eat and not drink caffeinated drinks for at least three hours before the test session as well as to refrain from intense physical exercise and alcohol consumption for 12 h before the experiment. Female participants completed the stress test during the luteal phase of their menstrual cycle to control for hormonal influences. The study protocol lasted approximately 90 min and comprised three sequential phases: a pre-stress phase, a stress phase, and a post-stress phase.

##### Psychosocial Stress Protocol

The stress condition was the Trier Social Stress Test (TSST), following the Kirschbaum et al. (1993) protocol. It consisted of three sections: preparation, interview, and a calculation task, with each section lasting 5 minutes.

##### Blood collection and sample processing

At each designated time point (T0–T4), blood was collected into 7.5 mL EDTA tubes (S-Monovette® 7.5 mL Z, Sarstedt, Nümbrecht, Germany). Samples were centrifuged at 1,610 × g for 10 min. Plasma was subsequently aliquoted and stored at −80 °C until further analysis.

##### Serum cortisol measurements

For serum cortisol measurements, blood samples were collected in serum monovettes (S-Monovette® 7.5 mL Z, Sarstedt, Nümbrecht, Germany). After blood collection, the monovettes were left at room temperature for 30 min to allow the blood to coagulate. The monovettes were then centrifuged at 2500 g for 10 min at 20°C, divided into aliquots and stored at −80°C. Serum cortisol concentrations were determined by single analysis and quantified using a commercially available enzyme-linked immunosorbent assay (ELISA) kit (IBL International GmbH, Germany). The detection limit of cortisol was 4.03–800 ng/mL with an analytical sensitivity of 3.79 ng/mL.

#### Cohort 2: Exercise Study

##### Ethical approval and study participants

The study was conducted between November 12, 2021, and January 18, 2022, at the Department of Obstetrics and Gynecology and the Institute of Sports Medicine, Prevention and Rehabilitation, Paracelsus Medical University Salzburg, Austria.

This study is part of a larger investigation that included both recreationally active participants and professional cyclists (see ClinicalTrials.gov: NCT05359744). This publication focuses on the results of the recreationally active participants (n=15).

Fifteen healthy recreationally active males [VO₂max 46,9 mL·kg⁻¹·min⁻¹ (SD = 5.12)] participated in a single-day interventional study. Participants were eligible if they were between 18 and 40 years of age and provided written informed consent to participate in the exercise intervention. All procedures were conducted in accordance with the Declaration of Helsinki. The study protocol was approved by the Ethics Committee of the State of Salzburg (EK No. 1132/2021) and registered at ClinicalTrials.gov (NCT05359744).

##### General study design

We conducted a single-center, non-blinded, non-randomized prospective study to investigate molecular changes during cardiopulmonary exercise testing (CPET) across varying workload (W) intensities on a cycle ergometer. All assessments were performed in the morning following an 8–12 h overnight fast. Baseline evaluations included anthropometric measurements (body mass [kg], height [cm], and body mass index [BMI; kg·m⁻²]), resting 12-lead electrocardiography (ECG), resting heart rate, peripheral oxygen saturation, resting blood pressure, and medical history assessment. In addition, transthoracic echocardiography was performed. Venous blood samples were obtained via an indwelling venous catheter inserted into an antecubital vein, which remained in situ throughout the entire testing protocol.

Blood samples were collected at baseline prior to exercise (T0), after completion of the aerobic warm-up phase (T1), immediately after the incremental test to volitional exhaustion (T3), and during recovery at 10 min (T4) and 30 min (T5) post-exercise.

##### Cardiopulmonary exercise testing (CPET)

Participants first completed a 15-min warm-up at a workload of 1 W·kg⁻¹. Predominantly aerobic energy metabolism during the warm-up was verified by spirometric assessment (stable respiratory quotient without an increase in CO₂ production) and blood lactate measurements, defined as no increase ≥ 0.5 mmol·L⁻¹ during the final 5 min of the warm-up (Biosen S-line Clinic, EKF Diagnostic GmbH, Magdeburg, Germany). After a 5-min resting period, maximal oxygen uptake (VO₂max) was determined using an incremental ramp protocol on an electromagnetically braked cycle ergometer (Lode Excalibur Sport, Lode B.V., Groningen, Netherlands) allowing for a linear increase in workload. Initial workload and ramp increments were individualized based on participants’ BMI, targeting test termination due to voluntary exhaustion within 8–12 min. The test was terminated upon volitional exhaustion. Pulmonary gas exchange and ventilation were measured breath-by-breath using a computerized metabolic cart (Metalyzer 3B, Cortex Biophysik GmbH, Leipzig, Germany). Throughout the entire protocol, participants were continuously monitored using a 12-lead ECG (ECGpro Version 5.10, AMEDTEC Medizintechnik Aue GmbH, Aue, Germany). Blood pressure was measured at time points T0 through T4. Blood samples for lactate analysis were collected at T0, at minute 7 of the warm-up, at T1, and at T3.

##### Blood collection and sample processing

At each designated time point (T0–T4), blood was collected into 7.5 mL lithium heparin tubes (Sarstedt, Nümbrecht, Germany; 11.608.001). Samples were centrifuged at 1,610 × g for 10 min. Plasma was subsequently aliquoted and stored at −80 °C until further analysis.

#### Cohort 3:Bungee study

It was a monocenter, non-blinded, non-randomized prospective trial. The intervention group (n = 22) performed a bungee jump, while the control group (n = 10) was not exposed to any experimental stress setup. Participants were recruited between May 2021 and January 2022. The study was conducted with healthy, male participants. For the present analyses, molecular profiling was performed exclusively in the intervention group, focusing on within-subject stress-induced changes.

##### Ethics

The study was carried out on the premises of the Telekom Baskets Bonn and at the Department of Psychiatry and Psychotherapy, University Hospital Bonn. The trial protocol was approved by the Ethics Committee of the University Hospital Bonn, Germany (lfd. Nr. 028/21; ClinicalTrials.gov identifier: NCT05144022). The study was conducted in accordance with the Declaration of Helsinki.

##### Participants

After potential participants expressed their interest in performing a bungee jump, they were contacted by the study team to evaluate possible participation. All participants were male and had no relevant previous illnesses. None had ever done a bungee jump before. Exclusion criteria included any neurological or psychiatric pre-existing conditions, regular medication (except for thyroid medication), pathological fear of heights, excessive alcohol consumption, smoking, illegal drug use, cardiovascular or pulmonary diseases, serious eye diseases, spine fractures, and surgeries within the last 4-6 months. The same inclusion and exclusion criteria applied to the control subjects.

##### Procedures

Intervention Group (Bungee Jump Group) (n = 22)

At the baseline appointments (and at every other blood draw), participants were asked not to eat or drink (except for water) for at least 12 hours. Participants were informed in advance (by e-mail and/or telephone) about the experimental setup. After arriving in the examination room, participants provided written informed consent and underwent further screening assessments for eligibility. This included verbal queries of inclusion and exclusion criteria, physical, neurological, and psychopathological examinations, and an electrocardiogram conducted by medical doctors. If any abnormalities were found, indicating the participant did not meet the inclusion criteria or met the exclusion criteria, they were excluded from the study. Participants also provided a urine sample, tested for illegal substances. If illegal substances were found, the participant was excluded from the study. If no reasons for exclusion were found, participants completed questionnaires on demographics and mental state, followed by the measurement of pulse and blood pressure and a venous blood sample (T0, baseline).

7-10 days after the baseline appointment, the bungee jump took place. Again, all participants were asked not to eat or drink (except for water) for at least 12 hours. In total, seven venous blood draws were taken on the day of the jump. The first blood draw (T1) occurred upon arrival at the experimental site, on average 48 minutes before the jump, followed by the second blood draw on the ground shortly on average 20 min before the jump (T2) and the third blood draw on average five min after the jump (T3). Additional blood draws were performed 20 minutes after the jump (T4), one hour after the jump (T5), two hours after the jump (T6), and four hours after the jump (T7). Participants fasted throughout this period. Between blood samples, participants were asked to avoid movement and stay at the experimental site. At selected blood draws, participants completed several psychological state questionnaires and cognitive tasks were performed at specified time points, which are not part of this manuscript.

For the bungee jump, participants were lifted on a platform to a height of 60 meters by a crane. The technical execution of the jump, including safety measures, was carried out by an external company specialized in bungee jumping.

The follow-up appointment (T8) took place 7-10 days after the day of the jump. Again, blood was taken for PBMC and plasma isolation, and participants completed the same questionnaires as at T0.

##### Blood collection and sample processing

Blood was collected in three 7.5 ml lithium heparin sampling tubes (Sarstedt, 11.608.001) and two 4.9 ml lithium heparin sampling tubes (Sarsted, 04.1939.001) at each timepoint. 7.5 ml lithium heparin sampling tubes were centrifuged at 1610 x g for 10 min. Afterwards plasma was collected and stored at −80 °C until it was further processed.

#### Single Stress Items questionnaire

Subjective stress, arousal and enthusiasm levels were assessed using a questionnaire specially designed for the study called “Single Stress Items”. In the questionnaire, participants were asked on the day of the jump to indicate their level of stress, enthusiasm and arousal on a scale from 0 to 100 as part of an open questionnaire.

#### Acute restraint stress experiment

##### Animals

C57BL/6JRj male mice were obtained from obtained from Janvier (France). Animals were housed in SealSafe PLUS ventilated cages (Tecniplast) in groups of 4–5 under a reversed 12-h light–dark cycle (lights off at 09:30 a.m.) with food and water ad libitum. The temperature in the housing facility was kept constant at 22 °C. All animals were acclimatized to the facility for at least two weeks before experiments. Age-matched mice (8–10 weeks old) were used to minimize age-related variability. The day before injections, mice were weighed and single-housed to reduce handling-related stress and to standardize conditions.

##### Injections and Stress paradigm

Mice (n = 20) were randomly assigned to one of four experimental groups. KH-103 stock solution was prepared by dissolving KH-103 powder in 40% Captisol (Selleck Chemicals) in DMSO. For injections, the stock was diluted to obtain a working solution containing 5% KH-103 in Captisol/DMSO, formulated into 20% Solutol (GLPBIO) in 0.9% sterile saline (Moltox) (w/v), yielding a final formulation with 5% DMSO. The mixture was vortexed thoroughly and sonicated in a heated water bath until a homogeneous, clear solution was obtained.

All animals received a subcutaneous injection of either KH-103 (44 mg/kg) or vehicle and were then left undisturbed in their single-housing cage for 2 h. Depending on group assignment, mice were subsequently exposed either to acute restraint stress or to a handling control condition. Acute restraint stress was performed by placing mice individually into 50 mL Falcon tubes with a large air hole for 90 min. The restraint tube was placed in the single-housing cage for the duration of stress. Handling controls were briefly handled and returned to their single-housing cage for the same duration. Following the 90-min manipulation, all mice remained undisturbed for an additional 150 min before euthanasia.

##### Serum collection

Trunk blood was collected immediately after euthanasia via cervical dislocation and decapitation. Samples were allowed to clot at room temperature for 30 min and subsequently centrifuged at 2,500 × g for 15 min at 4 °C. The resulting serum was carefully transferred into fresh microcentrifuge tubes and stored at −80 °C until further analysis.

#### Plasma Proteomics of human cohorts

##### Sample preparation for deep plasma proteomics analysis

For plasma proteomic profiling, a predefined subset of participants and timepoints was selected from each cohort to balance longitudinal coverage of the acute and delayed stress response with the analytical depth and throughput of deep plasma proteomics. Timepoints were chosen to capture baseline, peak and recovery phases of the respective stress paradigms. In the TSST cohort (Cohort 1), *n* = 20 participants per group were analyzed at timepoints T1, T8, and T10. In the exercise cohort (Cohort 2), *n* = 12 participants were analyzed at T0, T1, T2, T3, and T4. In the bungee cohort (Cohort 3), *n* = 16 participants were analyzed at T0, T2, T3, T5, and T7.

We utilized a nanoparticle-based enrichment strategy (Seer Proteograph^TM^ XT) (Blume et al., 2020) to overcome the high abundance problem of blood plasma samples and enable deep plasma-derived proteomes. From the collected EDTA (Cohort 1) or Lithium-Heparin blood plasma (Cohort 2 and 3), 240 µl of each sample was used for protein enrichment by protein corona formation around two nanoparticle mixtures (NP1 and NP2). Following, samples were tryptically digested, and peptides cleaned up, all carried out within a working day on a liquid handling system following the manufacturer’s protocol. Afterwards, peptide concentrations were determined (Thermo Pierce^TM^Fluorescence Quantitative Peptide Assay), and peptides were vacuum-dried completely within the collection plate overnight. On the next day, 200 ng of peptides were loaded onto Evosep Evotip Pure trap columns following the manufacturer’s protocol. Eleven synthetic peptides were spiked in as internal standards and at equimolar concentrations.

##### Liquid chromatography-tandem mass spectrometry

For the exercise and bungee jumping cohorts, proteomics samples were analyzed using a liquid chromatography-tandem mass spectrometry (LC-MS/MS) system comprising an Evosep One chromatographic system (Evosep) and a timsTOF Pro 2 mass spectrometer (Bruker Daltonics). Peptides were separated employing the pre-installed 40-sample-per-day whisper method with 31-minute chromatographic gradients. Separation occurred on an Aurora Elite CSI analytical column (Ionopticks), 15 cm in length with a 75 µm inner diameter, packed with 1.7 µm C18 particles. The LC system was directly coupled to the mass spectrometer, with eluting peptides transferred to the MS via a Captive Spray nano-electrospray ion source, operating at a constant spray voltage of 1.5 kV. Data were acquired in data-independent acquisition mode with parallel accumulation-serial fragmentation (diaPASEF). The dual TIMS analyzer ran at 100% duty cycle, utilising 100 ms for both accumulation and ramping times. Peptides were isolated within a mass-to-charge range of 350–1100 m/z and an ion mobility range of 0.63–1.45 Vs/cm². Sixteen py_DiAID-optimized PASEF windows with variable sizes were used for acquisition. Collisional energies were incrementally increased from 24 eV at 0.63 Vs/cm² to 49 eV at 1.45 Vs/cm².

For the TSST cohort, proteomics samples were analyzed using a liquid chromatography-tandem mass spectrometry (LC-MS/MS) system comprising an Evosep One chromatographic system (Evosep) and an Orbitrap Astral mass spectrometer (Thermo Fisher Scientific). Peptides were separated using the pre-installed 40-sample-per-day Whispher Zoom method with a 32.5-minute effective gradient and an Aurora Elite analytical column (Ionopticks, 15 cm length, 75 µm inner diameter, 1.7 µM C18 material. Eluting peptides were directly transferred to the MS via a Nanospray Flex ionization source, operated with a constant spray voltage of 1.8 kV. Data was acquired in data-independent acquisition mode. Full MS spectra were acquired with the Orbitrap detector operated at a resolution of 240,000. The AGC target was set to 300 % with a maximum injection time of 5 ms. The scan range was set to 380 – 980 m/z and a full MS acquisition event was triggered every 600 ms. Fragment MS spectra were recorded using the Astral detector. DIA windows were set to a width of 3 m/z with a total of 199 DIA windows. Fragmentation was achieved by Higer collisional dissociation (HCD) using relative HCD collision energies of 25%. The AGC target was set to 300 % with a maximum injection time of 5 ms.

##### Data analysis

Raw data files were processed by Spectronaut (version 19.1) with the Direct DIA search option using the built-in search engine Pulsar and the following default settings: the number of trypsin missed cleavages was set to 2, with minimum and maximum peptide lengths of 7 and 52 amino acids, respectively. PSM, peptide, and protein group identification FDR thresholds were set to 0.01. The search was performed using the reviewed UniProt database (H. sapiens, downloaded on 02/15/2023, SwissProt). Carbamidomethyl (C) was set as a fixed modification, and Oxidation (M) and Acetyl (Protein N-term) were set as variable modifications. IDPicker was used as Protein Inference Strategy. A proteotypicity filter was applied during analysis. Quantification of peptides was performed at the MS2 level.

##### Normalization, Filtering and Transformation

Bioinformatic analyses were conducted using R (v4.3.2). For the exercise and bungee jumping cohorts, batch effect normalization between plates was achieved by applying the ComBAT algorithm using the sva R package (3.50.0) (Leek et al., 2012). Before ComBAT, missing peptide intensities were imputed using a width of 0.3 and down-shift of 1.8, equivalent to the Perseus method, with a seed of 20241874. Imputed adjusted intensitiies were subsequently removed before directLFQ (0.2.18 with Anaconda Python 3.11.5) (Ammar et al., 2023) was used to quantify protein intensities based on adjusted peptide intensities.

For the TSST cohort, no systematic batch effects were detected based on exploratory data analysis and quality control metrics. Therefore, no additional batch correction was applied, and normalized protein intensities obtained from Spectronaut using the MaxLFQ algorithmwere used directly for downstream analyses

Protein intensities were log2-transformed and filtered to retain protein groups quantified in at least 50% of samples in at least one time point. No missing value imputation was performed.

##### Statistical Analysis

Plasma proteomics data were analyzed in R using the limma package (Ritchie et al., 2015). Longitudinal differential abundance analysis was conducted using linear models with empirical Bayes moderation to assess within-subject changes across time points. To account for repeated measurements within individuals, intra-subject correlation was estimated using the duplicateCorrelation function and incorporated into the linear modeling framework. Two complementary statistical approaches were applied: (i) moderated paired t-tests for selected pairwise comparisons between time points, and (ii) moderated F-tests to assess overall temporal effects across all time points. P values were adjusted for multiple testing using the Benjamini–Hochberg false discovery rate (FDR) procedure (Benjamini and Hochberg, 1995), and adjusted p values < 0.05 were considered statistically significant.

#### Hierarchical clustering and pathway enrichment analysis

Unsupervised clustering of significantly regulated proteins was performed in the primary analysis based on their longitudinal expression profiles using hierarchical clustering with Euclidean distance and complete linkage. These cluster assignments were subsequently reused for aggregated visualization.

For aggregated heatmaps, protein intensities were z-score standardized across all samples. Z-scores were averaged across participants for each time point to obtain mean temporal profiles. Proteins were ordered according to their original cluster assignments, and no additional clustering was performed. Heatmaps were generated using the pheatmap and ComplexHeatmap packages in R.

GO enrichment analysis was performed using the clusterProfiler R package (Yu et al., 2012). Enriched terms were identified with a q-value cutoff of 0.05, adjusted using the Benjamini-Hochberg method (Benjamini and Hochberg, 1995). Redundant GO terms were simplified using the simplify function with a similarity cutoff of 0.7.

#### Proteomics of mouse serum

Serum proteomics of the mouse cohort was performed using the same nanoparticle-based protein enrichment strategy as described above. Due to limited sample availability, 30 µl of serum was used per sample. Samples were analyzed by liquid chromatography–tandem mass spectrometry (LC-MS/MS) using an Evosep One chromatographic system (Evosep) coupled to an Orbitrap Astral mass spectrometer (Thermo Fisher Scientific), with instrument settings identical to those applied in the TSST cohort.

Database searching and protein quantification were performed in Spectronaut as described above, using the UniProt *Mus musculus* reference proteome (SwissProt, downloaded on 18/11/2021). Protein intensities were log2-transformed, and protein groups were retained if quantified in at least 50% of samples in at least one experimental group. No missing value imputation was performed.

Statistical analysis was carried out in R using the limma package. Moderated t-tests were applied for comparisons between groups. P-values were adjusted for multiple testing using the Benjamini–Hochberg false discovery rate (FDR) procedure; adjusted p-values < 0.05 were considered statistically significant.

#### Targeted Metabolomics of human cohorts

##### Sample preparation

Polar metabolite fractions were extracted from 50 µl of human plasma using methyl tert-butyl ether (MTBE):methanol:water (50:30:20, v/v/v, all LC-MS-grade). Medium blanks were processed in parallel. For 50 µl of human plasma, 1 ml pre-chilled (−20 °C) extraction buffer was added, samples were vortexed and kept on ice. Extracts were incubated in a cooled thermomixer (4 °C, 1,500 rpm, 30 min, Bioer), centrifuged (21,000 *× g*, 10 min, 4 °C, Eppendorf), and supernatants transferred. To induce phase separation, 200 µl MTBE (Sigma-Aldrich) and 150 µl water (Biosolve) were added and the mixture was incubated in a thermomixer (10 min, 15 °C, Bioer), followed by centrifugation (16,000 *× g*, 10 min, 15 °C, Eppendorf). The lipid (upper) and polar (lower) phases (∼650 µl and ∼600 µl, respectively) were collected, dried in a SpeedVac (room temperature, 4–6 h for polar fractions, Eppendorf), and stored at −80 °C.

##### LC-MS acquisition

Semi-quantitative determination of NADPH, NADP, NADH, NAD, AMP, ADP, ATP, Sedoheptulose-7-Phosphate, 6-Phosphogluconate, Fructose-6-Phosphate, Glucose-6-Phosphate, Pentose-Monophosphates (total), 3-Phospho-Glycerate, Phosphoenole-Pyruvate, Alpha-Ketoglutarate, Malate, Succinate, Fumarate, Lactate, Pyruvate, Glycolate, and Glyoxylate was performed using a LC-MS/MS. The chromatographic separation was performed on an Agilent Infinity II 1290 HPLC system using a SeQuant ZIC-pHILIC column (150 × 2.1 mm, 5 μm particle size, peek coated, Merck) connected to a guard column of similar specificity (20 × 2.1 mm, 5 μm particle size, Phenomoenex) a constant flow rate of 0.1 ml/min with mobile phase A comprised of 10 mM ammonium acetate in water, pH 9, supplemented with medronic acid to a final concentration of 5 μM and mobile phase B comprised of 10 mM ammonium acetate in 90:10 acetonitrile to water, pH 9 at 40° C.

The injection volume was 2 µl. The mobile phase profile consisted of the following steps and linear gradients: 0 – 1 min constant at 75 % B; 1 – 6 min from 75 to 40 % B; 6 to 9 min constant at 40 % B; 9 – 9.1 min from 40 to 75 % B; 9.1 to 20 min constant at 75 % B. An Agilent 6495 ion funnel mass spectrometer was used in positive and negative mode with an electrospray ionization source and the following conditions: ESI spray voltage 3000 V(−)/1500 V(+), nozzle voltage 2000 V(−)/1000 V(+), sheath gas 300° C at 9 l/min, nebulizer pressure 20 psig and drying gas 100° C at 11 l/min. Compounds were identified based on their mass transition and retention time compared to standards. Chromatograms were integrated using MassHunter software (Agilent, Santa Clara, CA, USA). Relative Abundance was determined based on peak area.

Quantitative determination of neurotransmitters was performed using a LC-MS/MS. The chromatographic separation was performed on an Agilent Infinity II 1290 HPLC system using a ZicHILIC SeQuant column (150 × 2.1 mm, 3.5 μm particle size, 100 Å pore size) connected to a ZicHILIC guard column (20 × 2.1 mm, 5 μm particle size) (Merck KgAA) a constant flow rate of 0.35 ml/min with mobile phase A being 0.1 % Formic acid in 99:1 water:acetonitrile (Honeywell, Morristown, New Jersey, USA) and phase B being 0.1 % formic acid 99:1 acetonitrile:water (Honeywell, Morristown, New Jersey, USA) at 25° C. The injection volume was 2 µl. The mobile phase profile consisted of the following steps and linear gradients: 0 – 8 min from 80 to 60 % B; 8 – 10 min from 60 to 10 % B; 10 – 12 min constant at 10 % B; 12 – 12.1 min from 10 to 80 % B; 12 to 15 min constant at 80 % B. An Agilent 6470 mass spectrometer was used in positive mode with an electrospray ionization source and the following conditions: ESI spray voltage 4500 V, nozzle voltage 1500 V, sheath gas 300° C at 12 l/min, nebulizer pressure 30 psig and drying gas 250° C at 11 l/min.

Compounds were identified based on their mass transition and retention time compared to standards. Chromatograms were integrated using MassHunter software (Agilent, Santa Clara, CA, USA). Relative abundance was determined based on the peak area.

Chromatograms were integrated using MassHunter software (Agilent, Santa Clara, CA, USA).

##### Statistical analysis

Metabolite intensities were log₂-transformed prior to analysis; zero values were treated as missing. Metabolites with a detection rate below 70% across all samples were excluded from analysis. Bungee-Jumping and Exercise cohorts: The effect of time on metabolite levels was assessed using linear mixed-effects models (LME) with timepoint as a fixed effect (factor) and participant as a random intercept to account for repeated measurements: log₂(intensity) ∼ factor(timepoint) + (1|participant). Model significance was evaluated by likelihood ratio test (LRT) comparing the full model against a null model without the timepoint term. P-values were adjusted for multiple comparisons using the Benjamini-Hochberg false discovery rate (FDR) procedure across all tested metabolites. Metabolites with FDR < 0.05 were considered statistically significant. Post-hoc pairwise comparisons against baseline (t = 0) were performed using the emmeans package (treatment vs. control contrasts) with Benjamini-Hochberg correction.

TSST cohort: Due to the small number of repeated measurements (3 timepoints) and moderate sample size (n = 20 per group), linear mixed-effects models showed singular fit (random intercept variance ≈ 0). Therefore, the non-parametric Friedman test (equivalent to one-way repeated measures ANOVA) was used to assess within-group time effects, separately for the control group, the PD group, and all participants combined. Post-hoc pairwise comparisons against baseline (t = −15 min) were performed using Wilcoxon signed-rank tests with Benjamini-Hochberg correction. All analyses were performed in R (version 4.4) using the packages lme4, emmeans, and base R.

#### Targeted metabolomics of steroid hormones

Steroids were determined by isotope dilution mass spectrometry (IDMS) using the MassChrom® Steroids assay for the quantitative determination of steroids in serum by liquid chromatography tandem mass spectrometry (LC-MS/MS) (Chromsystems GmbH, Gräfelfing/Germany).

Sample preparation was carried out with sample clean up columns included in the assay. In short: 500 µl of sample was mixed with 50 µl Internal Standard Mix and 450 µl Extracton Buffer in the equilibrated columns. The columns were centrifuged at 400 x g for one minute and the effluent discarded. After two wash steps steroids were eluted by 500 µl Elution buffer into autosampler vials. Eluents were dried at 50°C under nitrogen to dryness. The samples were reconstituted in 100 µl reconstitution buffer. 2 µl to 20 µl were injected into the mass spectrometry system dependent on the panel and included analytes. An AB Sciex API 5500 Qtrap LC/MS/MS (AB Sciex LLC, Framingham, MA, USA) equipped with an Agilent 1200 HPLC system (Agilent, Santa Clara, CA, USA) was used in multiple reaction mode (MRM). Aldosterone, Androstenedione, Cortisol, 11-Deoxycortisol, 21-Deoxycortisol, Cortisone, Corticosterone, 11-Deoxycorticosterone, Dehydroepiandrosterone (DHEA), Dehydroepiandrosterone sulfate (DHEAS), Progesterone, 17-OH-Progesterone, Testosterone and Dihydrotestosterone (DHT) were quantified in two different panels. Aldosterone and DHEAS were measured in negative, the other analytes in positive mode. A full calibration with seven calibration levels was carried out within each run. Three control samples with different levels were included, too. Data were evaluated by Analyst 1.7.2. using the matching internal standard.

Steroid concentrations below the lower limit of quantification (LOQ) were imputed as LOQ/2 prior to analysis. For each steroid, a linear mixed model (LMM) was fitted with time point as a fixed effect and a random intercept per subject to account for repeated measurements (R packages *lme4* and *lmerTest*). To control for multiple comparisons across the 14 measured steroids, p-values from the omnibus F-test (Type III) were adjusted using the Benjamini–Hochberg false discovery rate (FDR) procedure. For steroids surviving FDR correction (FDR < 0.05), post-hoc comparisons of estimated marginal means (EMMs) were conducted between each time point and baseline (BE1) using Dunnett-type contrasts, with Holm–Bonferroni correction applied within each steroid.

#### EV profiling by multiplexed bead-based flow cytometry

##### Quality control of plasma samples

Blood plasma dedicated for EV analysis was inspected for hemolysis and platelet background according to the Minimal Information for Blood EV research (MIBlood-EV) guidelines (Lucien et al., 2023) to assess the quality of blood plasma samples and identify potential confounders of EV analysis.

The presence of hemolysis was assessed qualitatively by visual inspection of plasma samples using the hemolysis reference palette (Kosecki et al., 2021). All plasma samples used for EV preparations in this study were below 100 mg/dl hemoglobin and thus suitable for further processing.

The residual platelet concentration in plasma samples was determined by flow cytometry. 20 µl of plasma were diluted with 80 µl of FACS staining buffer (0.5 % BSA, 2mM EDTA and 0.1 % NaN3 in PBS) and stained with 5 µl of CD61-FITC antibody (cat. no. 336404, Biolegend). Measurement was performed at an Attune Nxt flow cytometer (Thermo Fisher Scientific) to confirm minimal residual platelet concentration in plasma samples. The residual platelet concentration for all plasma samples was below 2×10^6^ platelets/ml.

##### EV separation from plasma samples by immuno-affinity capture

EVs were separated by immuno-affinity capture using the EV Isolation Kit CD63 (Miltenyi Biotec) according to the manufacturer’s instructions. Briefly, plasma was thawed in a water bath at 37 °C and diluted to a final volume of 2 ml in PBS. 50 µl of CD63 antibody-coupled magnetic beads were added to 2 ml of diluted plasma, following 1 hour of incubation at RT under constant shaking. Diluted samples were loaded onto µ columns attached to a μMACS™ Separator. CD63^+^ EVs were magnetically captured, washed, and eluted in 120 µl isolation buffer. 120 µl eluates were immediately used as input for the MACSPlex EV Kit IO, human, or MACSPlex EV Kit Neuro, human, platforms (Miltenyi Biotec, Bergisch Gladbach). Five subjects were processed with lower plasma input (350 µl instead of 600 µl), however normalized data were in the same range.

##### EV profiling by multiplexed bead-based flow cytometry

Phenotyping of CD63^+^EVs was performed by multiplexed bead-based flow cytometry as described (Brahmer et al., 2023) using the platforms MACSPlex EV Kit IO, human, and MACSPlex EV Kit Neuro, human, (Miltenyi Biotec). CD63^+^ EVs (120 µl) were incubated overnight with 15 µl of MACSPlex EV Kit IO, human, or MACSPlex EV Kit Neuro, human, capture beads (containing 39 different antibody-coated bead subsets). Samples and a buffer control were further processed according to the manufacturer’s instructions of the MACSPlex EV Kit, human (Miltenyi Biotec). Briefly, samples were washed and incubated for 1 h at RT with 15 µl of a cocktail of APC-labelled tetraspanin detection antibodies anti-CD9, anti-CD63 and anti-CD81. Flow cytometric analysis of EV samples was performed using the MACSQuant^®^ Analyzer 16 (Miltenyi Biotec) and the corresponding software MACSQuantify™ (Version 2.13.3). The signal intensity measured for each target was subtracted by the signal intensity for the respective target in the buffer control measurement. Negative values were set to zero. Signal intensities exceeding a predefined threshold with a value of 5, as specified by the manufacturer, were considered as robust and quantifiable signals.

#### EV isolation using membrane affinity method

EV isolation was performed at room temperature (RT) using the exoEasy Maxi Kit (QIAGEN), which is based on a membrane affinity method, following the manufacturer’s protocol. Briefly, 1 mL of plasma was thawed and centrifuged at 3000 × *g* for 5 min. The resulting supernatant was mixed with an equal volume of buffer XBP, loaded onto the exoEasy column, and centrifuged at 500 × *g* for 1 min. The flow-through was discarded, and the column membrane was washed with 10 mL of buffer XWP by centrifugation at 4900 × *g* for 5 min. Finally, EVs were eluted in 400 µL of buffer XE by centrifugation at 500 × *g* for 5 min. For the preparation of EV lysates, EV isolates were lysed in an appropriate volume of SDS lysis buffer (1% SDS, 50 mM Tris-HCl, pH 8), freshly supplemented with 1x cOmplete™ EDTA-free protease inhibitor cocktail (Roche, 4693132001) for 20 min on ice. Subsequently, samples were pulse-sonicated, vortexed and centrifuged for 10 min at 4°C at 13,000 rpm. The resulting supernatants were transferred to new 1.5 ml reaction tubes and analyzed by immunoblotting and LC-MS proteomics.

#### Proteomics of EVs

For proteomics of EVs, each sample was quantified using the Bicinchoninic acid assay (BCA Protein Assay Kit, Thermo Scientific, 23225). The aliquots to be quantified were diluted twenty times before adding them to a 96-well microplate (Thermo Scientific, 2205). Afterwards the working solution was prepared by mixing reagent A and reagent B with ratio A:B in a ratio of 50:1. 200 µL of the working solution was then added, and the microplate was incubated at 37 °C for 30 min. Finally, the microplate was quantified using SPARK® Multimode Microplate Reader and SPARKControl Magellan analysis software. For each sample, 10 µg of protein was mixed with reduction/alkylation buffer to achieve final concentrations of 1% SDS, 10 mM TCEP (tris(2-carboxyethyl)phosphine), and 30 mM CAA (chloroacetamide) in a total volume of 50 µL. Samples were heated at 90°C for 10 minutes to ensure complete denaturation, reduction, and alkylation. MagReSyn Hydroxyl microparticles (ReSyn Biosciences) were used at a 1:4 protein-to-bead ratio (w/w) and equilibrated twice with 70% acetonitrile prior to use. The reduced and alkylated samples were added to the equilibrated microparticles, and acetonitrile was added to a final concentration of 70% to induce on-bead protein aggregation. Following a 10-minute incubation, magnetic separation was performed.

Beads were washed sequentially with 100% acetonitrile and 70% ethanol to remove SDS and other contaminants. Subsequently, beads were resuspended in digestion buffer (50 mM TEAB, pH 8.5) and proteins were enzymatically digested using a Trypsin/Lys-C mixture overnight at 37°C. Digestion was quenched by acidification with TFA to a final concentration of 1% and loaded onto Evotips (Evosep) for subsequent LC-MS/MS analysis. LC-MS/MS measurements and statistical analysis were performed using a liquid chromatography-tandem mass spectrometry (LC-MS/MS) system comprising an Evosep One chromatographic system (Evosep) and a timsTOF Pro 2 mass spectrometer (Bruker Daltonics) as described previously (see section Plasma Proteomics of human cohorts)

#### Capillary Immunoblotting

For capillary immunoblotting, plasma samples were diluted with Sample Buffer (ProteinSimple, # 042-195) in ratio 1:1, mixed with a master mix containing dithiothreitol (DTT) and fluorescent molecular weight marker (EZ Standard Pack, 12–230 kDa; ProteinSimple), and denaturated at 95°C for 5 minutes. Proteins were separated and analyzed by capillary electrophoresis on Jess^TM^ (ProteinSimple; San Jose, CA, United States) according to manufacturer’s protocol using the 12–230 kDa cartridges. The following primary antibodies were utilized for immunodetection: rabbit anti-LC3B (1:50, Cell Signaling Technology, #2775), rabbit HnrnpD (1:50, Cell Signaling Technology, #12382) and rabbit CD9 (1:50, Cell Signaling Technology, #D8O1A).

Anti-Rabbit detection module (ProteinSimple, San Jose, CA, United States) and anti-mouse detection module (ProteinSimple, San Jose, CA, United States) were used for detecting primary antibodies.

#### PBMC Stimulation experiment

##### Ethics and PBMC isolation

The overall study design is described elsewhere (Cohort#4-Stress Less Study) (Mackert et al., 2024). The study was carried out at the Department of Psychiatry and Psychotherapy, University Hospital Bonn, and the trial protocol was approved by the Ethics Committee of the University Hospital Bonn in Germany (Venusberg Campus 1, 53127 Bonn, ID:319/20). Additionally, the study was registered with ClinicalTrials.gov (ClinicalTrials identifier: NCT04823806). For the isolation of PBMCs, we took blood draws from six healthy male participants at baseline (without any supplementation started). Blood was collected in three 7.5 ml lithium heparin sampling tubes (Sarstedt, 11.608.001). 7.5 ml lithium heparin sampling tubes were centrifuged at 1610 x g for 10 min. The lithium heparin tubes were carefully loaded on Pancoll solution (PAN Biotech, P04-60500) in Leucosep tubes (Greiner BioOne, 2007473/2026886) and centrifuged at 684 x g for 30 min (brakeless running down). PBMCs were enriched by selecting the interphase of the Pancoll gradient. PBMCs of the interphase were washed three times with ice-cold PBS.

##### Stimulation

PBMCs were isolated from 6 donors, each contributing 3 × 7.5 mL Li-heparin blood collection tubes (Sarstedt, 11.608.001), according to the standard PBMC isolation protocol described above. Isolated cells were counted and initially cultured in complete RPMI growth medium (Thermo Fisher Scientific, 61870010) containing 10 % FBS (PAN Biotech, P30-3306) in 10 cm dishes. Prior to stimulation, cells were washed three times with RPMI without FBS, supplemented with NEAA (Thermo Fisher Scientific, 11140050) (stimulation medium), and seeded into 6-well plates in stimulation medium. Cells were stimulated with lactate (Sigma Aldrich, 71718, 15 mM,), β-hydroxybutyrate (BHB, 5 mM), lactoyl-phenylalanine (Lac-Phe, 50 µM), cortisol (Sigma-Aldrich, H0888, CORT, 150 nM), or vehicle control. After 1 h of incubation, supernatants were harvested, centrifuged at 300 × g, transferred into fresh tubes, and stored at −80°C until downstream analysis.

##### Proteomics of supernatants

Sample preparation for mass spectrometry analysis was performed using Protein Aggregation Capture (PAC) according to Batth et al. (2019), adapted for automated processing on an Agilent Bravo Liquid Handling System in 96-well plate format.

For each sample, 25 µL of supernatant in 1% SDS, 50 mM Tris-HCl was used as input. Samples were adjusted to a final volume of 50 µL containing 10 mM TCEP (tris(2-carboxyethyl)phosphine) and 30 mM CAA (chloroacetamide) for simultaneous reduction and alkylation. Samples were heated at 90°C for 10 minutes to ensure complete denaturation, reduction, and alkylation.

MagReSyn Hydroxyl microparticles (ReSyn Biosciences) were used and equilibrated twice with 70% acetonitrile prior to use. The reduced and alkylated samples were added to the equilibrated microparticles, and acetonitrile was added to a final concentration of 70% to induce on-bead protein aggregation. Following a 10-minute incubation, magnetic separation was performed.

Beads were washed sequentially with 100% acetonitrile and 70% ethanol to remove SDS and other contaminants. Subsequently, beads were resuspended in digestion buffer (50 mM TEAB, pH 8.5) and proteins were enzymatically digested using a Trypsin/Lys-C mixture overnight at 37°C. Digestion was quenched by acidification with TFA to a final concentration of 1% and loaded onto Evotips (Evosep) for subsequent LC-MS/MS analysis.

LC-MS/MS measurements were performed using a liquid chromatography-tandem mass spectrometry (LC-MS/MS) system comprising an Evosep One chromatographic system (Evosep) and an Orbitrap Astral as described previously, but with slight variations (see section Plasma Proteomics of human cohorts). In total, 300 DIA fragment spectra were acquired with a window width of 2 m/z and a maximum injection time of 3 ms. All other parameters remained unchanged.

Database searching and protein quantification were performed in Spectronaut as described above, using the UniProt *Homo sapiens* reference proteome (SwissProt). Protein intensities were log2-transformed, and protein groups were retained if quantified in at least 50% of samples in at least one experimental group. No missing value imputation was performed.

Statistical analysis was carried out in R using the limma package. Moderated t-tests were applied for comparisons between groups. P-values were adjusted for multiple testing using the Benjamini–Hochberg false discovery rate (FDR) procedure; adjusted p-values < 0.05 were considered statistically significant.

#### Statistics and reproducibility

For data processing, Rstudio along with R language (v4.3.2 or v.4.1., Rcore team, 2019) and GraphPad Prism (versions 9.0 or 10.0) were employed.

Unless stated otherwise in the methods section of the respective measurement, missing values in repeated measures designs were analyzed using a mixed-effects model as implemented in GraphPad Prism. This mixed model uses a compound symmetry covariance matrix, and is fit using Restricted Maximum Likelihood (REML). In the absence of missing values, this method gives the same P values and multiple comparisons tests as repeated measures ANOVA. In the presence of missing values (missing completely at random), the results can be interpreted like repeated measures ANOVA.

## Data Availability

The mass spectrometry proteomics data will be deposited to the ProteomeXchange Consortium via the PRIDE partner repository and made publicly available upon publication. Confidential access for reviewers can be provided upon request during peer review.

All supplementary tables can be found at https://doi.org/10.5281/zenodo.20381885

## Code Availability

Analyses were performed using standard statistical libraries in R, as described in the Materials and Methods section. No custom software was developed for this study.

## Acknowledgements

We thank all lab members for suggestions and comments on the experiments and manuscript. We are grateful for the technical assistance provided by T. Schnitzler, C. Strauch-Heinz, M. Ebert and A. Gassen.

## Author Contributions

Conceptualization: TE, NCG

Methodology: TE, TB, JS, BH, JE, HS, EMK-A, NP, MS, FM, NCG

Software: TE, WB, SB, HH, JC

Validation: TE, TB, WB, JE, SK, EMK-A, NCG

Formal Analysis: TE, TB, WB, JE, SB, BS, FM, JC, NCG

Investigation: TE, TB, JS, BH, JE, SK, WB, EJ, MLe, DEH, HH, MCS, PB, MB, NB, TF, EH, MH, HJ-B, SK-S, VK, MK, EK, AK, MLa, KL, CL-F, BL, AL, AR, SL, SM, MM, KP, JT, ET, AZ, HS, AP, MBM, BS, BM, JC, KG, EMK-A, NP, MS, FM, NCG

Resources: AP, BS, JC, FM, EMK-A, NCG

Data Curation: TE, TB, WB, JE, EK, SK, BS, JC, EMK-A, FM, NCG

Writing - Original Draft: TE, TB, NCG Writing - Review & Editing: all authors Visualization: TE, TB, NCG

Supervision: TE, JC, PK, EMK-A, NP, FM, NCG

Project administration: TE, NP, FM, NCG

Funding acquisition: TE, FM, MBM, BJ, EMK-A, NCG

## Supplementary Data

All supplementary tables can be found at https://doi.org/10.5281/zenodo.20381885

Table S1 - Demographic information of TSST cohort

Table S2 - Moderated t test results of the combined TSST cohort

Table S3 - Moderated t test results of both groups of the TSST cohort separately

Table S4 - GO analysis results oft he TSST cohort

Table S5 - Moderated t test results comparing both groups in the TSST cohort

Table S6 - Results from interaction test between PD patients and controls

Table S7 - Log2-transformed protein intensities of plasma proteomics from the TSST cohort

Table S8 - Demographic information of exercise cohort

Table S9 - Log2-transformed protein intensities of plasma proteomics of the exercise cohort

Table S10 - Moderated t test and Ftest results of the exercise cohort, including cluster annotations

Table S11 - GO analysis results of the exercise cohort

Table S12 - Demographic information of the bungee jump cohort

Table S13 - Log2-transformed protein intensities of plasma proteomics of the bungee jump cohort

Table S14 - Moderated t test and Ftest results of the bungee cohort, including cluster annotations

Table S15 - GO analysis results of the bungee jump cohort

Table S16 - List of stress core proteins

Table S17 - Metabolomics results

Table S18 - Moderated t test results of EV proteomics of the bungee jump cohort

Table S19 - List of overlapping proteins between EVs and plasma

Table S20 - Log2-transformed protein intensities and moderated t test results of secretome proteomics

Table S21 - Log2-transformed protein intensities and moderated t test results of plasma proteomics from mice treated with GR-Protac or vehicle

